# North Andean origin and diversification of the largest ithomiine butterfly genus

**DOI:** 10.1101/105882

**Authors:** Donna Lisa De-Silva, Luísa L. Mota, Nicolas Chazot, Ricardo Mallarino, Karina L. Silva-Brandão, Luz Miryam Gómez Piñerez, André V.L. Freitas, Gerardo Lamas, Mathieu Joron, James Mallet, Carlos E. Giraldo, Sandra Uribe, Tiina Särkinen, Sandra Knapp, Chris D. Jiggins, Keith R. Willmott, Marianne Elias

## Abstract

The Neotropics harbour the most diverse flora and fauna on Earth. The Andes are a major centre of diversification and source of diversity for adjacent areas in plants and vertebrates, but studies on insects remain scarce, even though they constitute the largest fraction of terrestrial biodiversity. Here, we combine molecular and morphological characters to generate a dated phylogeny of the butterfly genus *Pteronymia* (Nymphalidae: Danainae), which we use to infer spatial, elevational and temporal diversification patterns. We first propose six taxonomic changes that raise the generic species total to 53, making *Pteronymia* the most diverse genus of the tribe Ithomiini. Our biogeographic reconstruction shows that *Pteronymia* originated in the Northern Andes, where it diversified extensively. Some lineages colonized lowlands and adjacent montane areas, but diversification here remained scarce. The recent colonization of lowland areas was reflected by an increase in the rate of evolution of species elevational ranges towards present. By contrast, speciation rate decelerated with time, with no extinction. The geological history of the Andes and adjacent regions have likely contributed to *Pteronymia* diversification by providing compartmentalized habitats and an array of biotic and abiotic conditions, and by limiting dispersal between some areas while promoting interchange across others.

## Introduction

The Neotropical region is the most biologically diverse area on Earth for most organisms and numerous studies have identified the world’s longest mountain range, the Andes, as a major centre of biodiversity^1^ and source for adjacent areas in groups as diverse as birds^2^, reptiles^3^, insects^4–6^ and plants^7,8^. The Andes, has been proposed as a major driver of diversification^9^. For instance, studies of some Andean plants have found some of the fastest diversification events reported, such as in the Andean Bellflowers, whose 550 species arose in the last 5 million years^8^. The Andes may have affected diversification rates in different ways, by offering scope for vicariant speciation due to the complex and intricate topography of the mountains, as well as by providing a large array of new environmental niches, thereby promoting adaptive speciation.

The timing of diversification within the Neotropical region is keenly debated and is linked to competing hypotheses about which biogeographic events have primarily driven speciation, extinction and dispersal events during the Cenozoic. However, the rate and geographic extent of surface uplift in the Andes is contentious, having varied through time and among different cordilleras^10,11^, and assessing the role of the Andes and the different phases of uplift on the timing of diversification is complex. The closure of the Panamanian Isthumus during the last 5 million years may have allowed substantial biotic interchange between Central and South America (but see^12^) or the Pebas system, a large network of shallow lakes and wetlands, which occupied the upper Amazon region, may have constrained dispersal and promoted local diversification until its drainage around 10-7 million years ago^13,14^. It is also suggested that climatic instability during the Pleistocene drove the diversification of extant Neotropical species, however, the importance of this mechanism remains controversial^9,15–19^.

Insects represent the largest fraction of terrestrial biodiversity but analyses of insect diversification remain scarce compared to other groups such as vertebrates and plants. Butterflies are one of the best studied insect groups and around 7800 Neotropical species have so far been documented^20^. Over the last decade, an increasing number of studies have used molecular phylogenetic trees to investigate the timing and mode of diversification in a variety of butterfly groups. Such works provide scope for comparative approaches that can help decipher whether common drivers or distinct processes have shaped patterns of diversification. They also allow us to assess the extent to which inferred patterns of diversification match those found in other organisms, particularly well-studied plants^21–23^ and birds^2,24^. Many butterfly studies indicate an important role for the Andes in the origin and diversification of new species. In some groups, diversification occurs mostly within the same elevational range, consistent with adaptation onto new resources (e.g., *Hypanartia*^25^, *Lymanopoda*^26^, some ithomiine butterflies^6,27^), whereas others show speciation across the elevational gradient (e.g., *Ithomiola*^28^). Even within some predominantly lowland groups the Andes have apparently played an important role, causing diversification contemporaneous with the Andean uplift and consequent major changes in climate and geography in the Neotropical region (e.g., *Taygetis*^29^, *Dione*^30^, *Heliconius*^31^). By contrast, the Andes seem to have had a limited impact on the diversification of some other groups, such as neotropical Troidini^32^

The Neotropical butterfly tribe Ithomiini (Nymphalidae: Danainae) is a diverse group with ca. 48 genera and 390 species, which are ubiquitous in humid forest throughout the Neotropical region from sea level to around 3000 m elevation. Commonly known as the clearwing butterflies because of the transparent wings in many species, they are well-known because of their involvement in Mullerian mimicry rings, whereby co-occurring unpalatable species converge in wing colour pattern amongst themselves, other Lepidoptera and some other insects^33^. Their diversity and broad distribution makes them a relevant study system to investigate patterns of spatial and temporal diversification in the Neotropics and they have previously been the focus of a number of diversification studies^4,6,34,35^. Most ithomiine clades have been found to show a peak of species richness in the Andes^36^, but the biogeographic histories that have led to this pattern are surprisingly diverse. The genus *Napeogenes* originated in the Andes and subsequently dispersed out of the mountains into the Amazon Basin^6^, whereas rates of colonization into the Andes from adjacent areas were found to be higher in the subtribe Godyridina^34^. The genus *Oleria* contains two main subclades, one of which diversified mainly in lowland Amazonian forests while the other diversified almost exclusively in high elevation Andean cloud forests^4^.

The genus *Pteronymia* Butler & Druce 1872, which belongs to the largest ithomiine subtribe, the Dircennina, is one of the most speciose ithomiine genera. The genus contains some 50 species (Lamas, 2004), although the species taxonomy has been highly confused in the past and is undergoing revision. *Pteronymia* butterflies occur throughout the Neotropics, with the most diverse communities found in east Andean cloud forests^27^. As part of our on-going effort to document patterns of spatial and temporal diversification of butterflies in the Neotropics, here we reconstruct a comprehensive, time calibrated phylogeny for the *Pteronymia* using multi-locus molecular data and morphological characters. We first assess whether *Pteronymia* is monophyletic and whether do species boundaries require re-definition. We then time-calibrate the phylogeny using a combination of larval host plant ages (Solanaceae) as maximal age constraints^37^ and estimates of divergence time between nymphalid butterfly genera from Wahlberg et al. (2009)^38^ to investigate biogeographical of diversification of *Pteronymia* in relation with the Andean uplift. Specifically, we addressed the following questions: (1) Have the Andes acted as a centre of origin and a centre of diversification for the genus *Pteronymia*? (2) Has there been many interchanges between the different regions of the Neotropics, particularly between the Andes and other regions, and between the central and northern Andes? (3) How has the elevational niche of *Pteronymia* species evolved through time, and is there evidence for adaptive radiation across elevation ranges? (4) How has *Pteronymia* diversified through time?

## Results

We obtained sequence data for a total of 166 *Pteronymia* specimens, representing 41 of the species recognized prior to our revision, and 47 of the species recognized after our revision (Table 1; see Supplementary Table S1 online). Species with no molecular data were *P. alcmena, P. alicia, P. calgiria, P. fumida, P. glauca* and *P. peteri*. A total of 87 morphological characters were coded for 52 species (after taxonomic revision, see below).

**Table 1.**
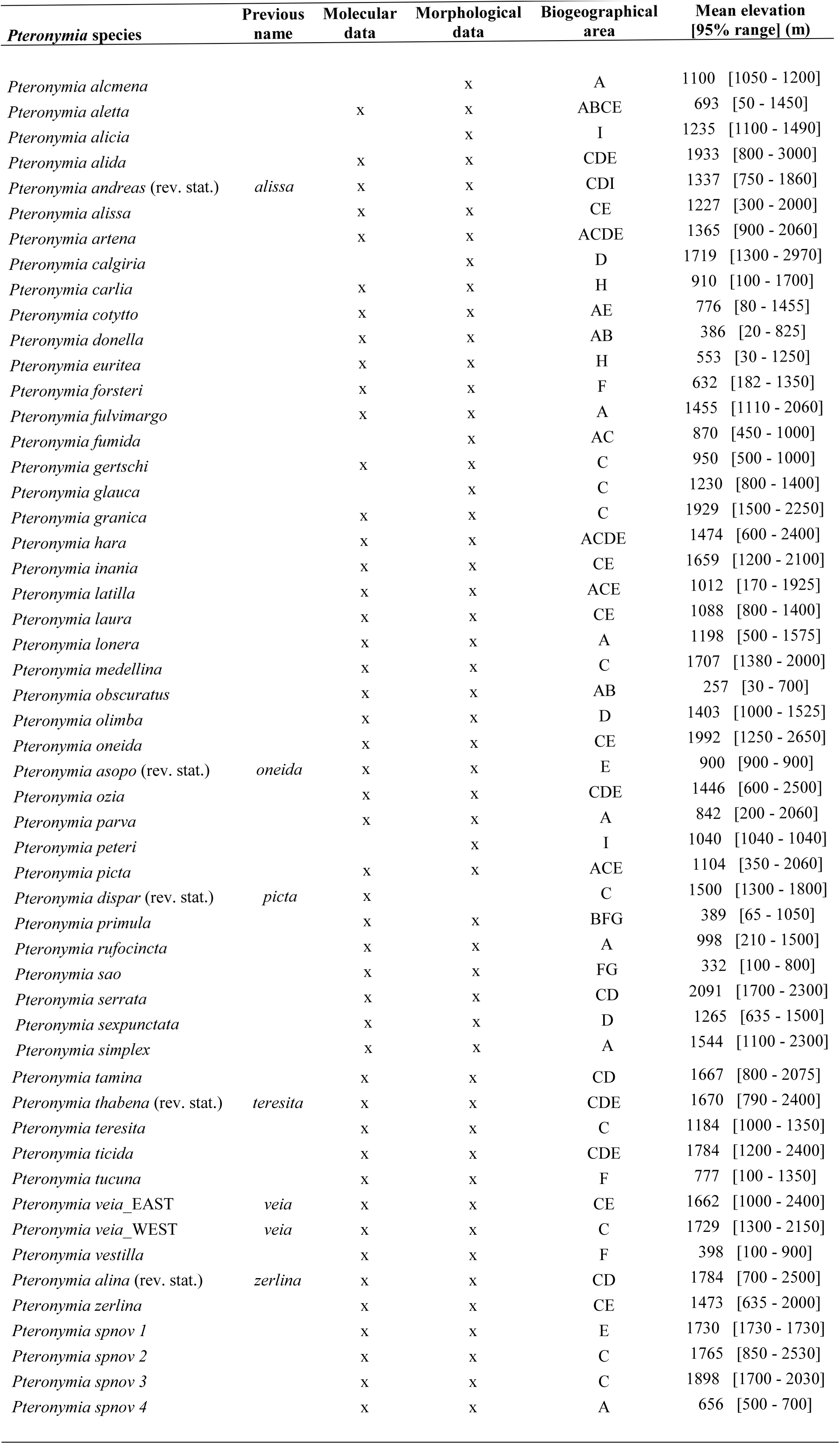
List of the species in the genus *Pteronymia*, including revised status. Availability of molecular and morphological data, distribution area and elevation mean and 95% range are reported for each species.

| *Pteronymia* species | Previous name | Molecular data | Morphological data | Biogeographical area | Mean elevation [95% range] (m) |
| --- | --- | --- | --- | --- | --- |
| *Pteronymia alcmena* |  |  | x | A | 1100 [1050 - 1200] |
| *Pteronymia aletta* |  | x | x | ABCE | 693 [50 - 1450] |
| *Pteronymia alicia* |  |  | x | I | 1235 [1100 - 1490] |
| *Pteronymia alida* |  | x | x | CDE | 1933 [800 - 3000] |
| *Pteronymia andreas* (rev. stat.) | *alissa* | x | x | CDI | 1337 [750 - 1860] |
| *Pteronymia alissa* |  | x | x | CE | 1227 [300 - 2000] |
| *Pteronymia artena* |  | x | x | ACDE | 1365 [900 - 2060] |
| *Pteronymia calgiria* |  |  | x | D | 1719 [1300 - 2970] |
| *Pteronymia carlia* |  | x | x | H | 910 [100 - 1700] |
| *Pteronymia cotytto* |  | x | x | AE | 776 [80 - 1455] |
| *Pteronymia donella* |  | x | x | AB | 386 [20 - 825] |
| *Pteronymia euritea* |  | x | x | H | 553 [30 - 1250] |
| *Pteronymia forsteri* |  | x | x | F | 632 [182 - 1350] |
| *Pteronymia fulvimargo* |  | x | x | A | 1455 [1110 - 2060] |
| *Pteronymia fumida* |  |  | x | AC | 870 [450 - 1000] |
| *Pteronymia gertschi* |  | x | x | C | 950 [500 - 1000] |
| *Pteronymia glauca* |  |  | x | C | 1230 [800 - 1400] |
| *Pteronymia granica* |  | x | x | C | 1929 [1500 - 2250] |
| *Pteronymia hara* |  | x | x | ACDE | 1474 [600 - 2400] |
| *Pteronymia inania* |  | x | x | CE | 1659 [1200 - 2100] |
| *Pteronymia latilla* |  | x | x | ACE | 1012 [170 - 1925] |
| *Pteronymia laura* |  | x | x | CE | 1088 [800 - 1400] |
| *Pteronymia lonera* |  | x | x | A | 1198 [500 - 1575] |
| *Pteronymia medellina* |  | x | x | C | 1707 [1380 - 2000] |
| *Pteronymia obscuratus* |  | x | x | AB | 257 [30 - 700] |
| *Pteronymia olimba* |  | x | x | D | 1403 [1000 - 1525] |
| *Pteronymia oneida* |  | x | x | CE | 1992 [1250 - 2650] |
| *Pteronymia asopo* (rev. stat.) | *oneida* | x | x | E | 900 [900 - 900] |
| *Pteronymia ozia* |  | x | x | CDE | 1446 [600 - 2500] |
| *Pteronymia parva* |  | x | x | A | 842 [200 - 2060] |
| *Pteronymia peteri* |  |  | x | I | 1040 [1040 - 1040] |
| *Pteronymia picta* |  | x | x | ACE | 1104 [350 - 2060] |
| *Pteronymia dispar* (rev. stat.) | *picta* | x |  | C | 1500 [1300 - 1800] |
| *Pteronymia primula* |  | x | x | BFG | 389 [65 - 1050] |
| *Pteronymia rufocincta* |  | x | x | A | 998 [210 - 1500] |
| *Pteronymia sao* |  | x | x | FG | 332 [100 - 800] |
| *Pteronymia serrata* |  | x | x | CD | 2091 [1700 - 2300] |
| *Pteronymia sexpunctata* |  | x | x | D | 1265 [635 - 1500] |
| *Pteronymia simplex* |  | x | x | A | 1544 [1100 - 2300] |

|  |  |  |  |  |  |  |
| --- | --- | --- | --- | --- | --- | --- |
| *Pteronymia tamina* |  | x | x | CD | 1667 | [800 - 2075] |
| *Pteronymia thabena* (rev. stat.) | *teresita* | x | x | CDE | 1670 | [790 - 2400] |
| *Pteronymia teresita* |  | x | x | C | 1184 | [1000 - 1350] |
| *Pteronymia ticida* |  | x | x | CDE | 1784 | [1200 - 2400] |
| *Pteronymia tucuna* |  | x | x | F | 777 | [100 - 1350] |
| *Pteronymia veia*_EAST | *veia* | x | x | CE | 1662 | [1000 - 2400] |
| *Pteronymia veia*_WEST | *veia* | x | x | C | 1729 | [1300 - 2150] |
| *Pteronymia vestilla* |  | x | x | F | 398 | [100 - 900] |
| *Pteronymia alina* (rev. stat.) | *zerlina* | x | x | CD | 1784 | [700 - 2500] |
| *Pteronymia zerlina* |  | x | x | CE | 1473 | [635 - 2000] |
| *Pteronymia spnov 1* |  | x | x | E | 1730 | [1730 - 1730] |
| *Pteronymia spnov 2* |  | x | x | C | 1765 | [850 - 2530] |
| *Pteronymia spnov 3* |  | x | x | C | 1898 | [1700 - 2030] |
| *Pteronymia spnov 4* |  | x | x | A | 656 | [500 - 700] |

### Taxonomy

Molecular phylogenies of all *Pteronymia* specimens generated by maximum likelihood and Bayesian inference were largely congruent (see Supplementary Figs. S1-S2 online), and showed that the genus *Pteronymia* is monophyletic. Several internal nodes had moderate to low support.

The molecular data suggested that several changes to the species-level taxonomy were warranted. Four cases concern species occurring on both slopes of the Andes, where the molecular data suggest that east and west Andean subspecies are not sister taxa, instead grouping with other related species. In all cases there are no genitalic characters that exclusively support the former classification, which was instead based on attempts to group similar allopatric phenotypes as subspecies of more widespread species^20^. We therefore split each of the original four species into east and west Andean species.

#### Pteronymia zerlina

Hewitson, 1856. This species formerly included taxa from both east and west of the Andes^20,39^, ranging from Venezuela to western Ecuador and Bolivia. Sequenced material comes from eastern and western Ecuador, representing the taxa formerly known as *P. zerlina pronuba* Hewitson, 1870 (west) and *P. zerlina machay* T. & L. Racheli, 2003 (east). *Pteronymia zerlina zerlina* Hewitson, 1856 was described from ‘New Granada’ and the original illustrations and type material in the BMNH match specimens from the Cauca valley in west Colombia. Their smaller size and broad white forewing translucent band are similar to *P. zerlina pronuba* and it thus seems likely that these two taxa are conspecific. In the eastern Andes, populations throughout Peru apparently link east Ecuadorian *P. zerlina machay* with the Bolivian *P. zerlina alina* Haensch, 1909, and thus we treat *P. alina* as a distinct species (**rev. stat.**) and transfer to it the following Peruvian and Ecuadorian taxa: *P. a. machay*, *P. a. mielkei* Lamas, 2003 (**rev. stat.**). The status of Venezuelan and central Colombian populations is currently unknown, so for the moment we retain them in *P. zerlina.* The molecular results are consistent with significant differences in the immature stages of *P. zerlina zerlina* and *P. zerlina machay* (now *P. alina machay*) described by^40^, who also suggested the likelihood that these taxa were not conspecific.

#### Pteronymia veia

Hewitson, 1853. This species formerly included taxa from both east and west of the Andes^20,39^, ranging from Venezuela to western Ecuador and northeastern Peru. Sequenced taxa include an undescribed taxon from western Ecuador and *P. veia linzera* Herrich-Schäffer, 1865 from eastern Ecuador. As with *P. zerlina*, the status of Venezuelan and Colombian populations is unknown, but unfortunately there are no described west Colombian taxa which can be reliably associated with the undescribed west Ecuadorian taxon. We treated east and west Ecuadorian taxa as distinct species, but for the moment do not make any nomenclatural changes.

#### Pteronymia alissa

Hewitson, 1869. This species formerly included taxa from both east and west ofthe Andes^20,39^, ranging from Venezuela to western Ecuador and Bolivia. Sequenced taxa include the the nominate subspecies from western Ecuador, and *P. alissa andreas* Weeks, 1901 from eastern Ecuador. The status of Venezuelan taxa is currently uncertain, so for the moment we retain them in *P. alissa* and just separate the east Andean *P. andreas* (**rev. stat.**) as a separate species, except for the subspecies *dorothyae*, which clusters with *P. andreas* in the phylogenetic trees (see Supplementary Figs. S1-S2 online).

#### Pteronymia teresita

Hewitson, 1863. This species formerly included taxa from both east and west ofthe Andes^20,39^, occurring in western Ecuador and from eastern Colombia to Bolivia. Sequenced taxa include west Ecuadorian *P. teresita teresita* and east Ecuadorian *P. teresita thabena* (Hewitson, 1869). We here separate out east Andean populations as a separate species, which both share a distinctive female with colorless translucent hindwing and yellow translucent forewing, including the following: *P. thabena thabena*, *P. thabena denticulata* Haensch, 1905 (**rev. stat.**).

#### Pteronymia oneida

Hewitson, 1855. This species formerly included taxa from western Colombia to Venezuela and along the eastern Andes to northern Peru^20,39^. *Pteronymia oneida asopo* (C. & R. Felder, 1865) occurs in the Cordillera de la Costa in northern Venezuela, and appeared distantly related to east Ecuadorian *P. oneida oneida* in the molecular tree. As with related species (*P. zerlina*, *P. veia*), there are no genitalic characters that group *asopo* with *oneida*, and we therefore treat it as a distinct species, *P. asopo* (**rev. stat.**). The status of several other recently described^39^ Venezuelan taxa of *P. oneida* remains to be determined, and for the moment they are retained in *P. oneida*.

#### Pteronymia picta

Salvin, 1869. This species formerly included taxa ranging from Costa Rica to central and western Colombia. Specimen locality data from Colombia are too imprecise and unreliable to confirm whether the Colombian *P. picta dispar* Haensch, 1905, apparently restricted to the northern Cordilleras Occidental and Central, is sympatric or not with *P. picta picta*, the range of which appears to broadly encompass that of *dispar* in Colombia. The molecular data suggest that *P. dispar* (**rev. stat**.) is not closely related to *P. picta picta* and Central American *P. picta notilla* Butler & H. Druce, 1872.

After our taxonomic changes the genus *Pteronymia* now comprises 53 species, making it the most diverse ithomiine genus.

### Morphological phylogeny

We performed a cladistic analysis of 87 adult and larval morphological traits, which resulted in a relatively poorly resolved tree, within which only several clades received moderate to strong support (see Supplementary Fig. S3 online). Notable clades include one containing six typically rare Andean species (*alida* clade: *P. alida*, *P.* sp. nov. 3, *P. inania*, *P. lonera*, *P. teresita* and *P. thabena*), supported by a number of genitalic characters. The immature stages (i. e., larvae and pupae) of this clade are also remarkably different in coloration and morphology from those of other *Pteronymia* species, and the distinctiveness of the genitalia and immature stages previously led to the description of a new genus, *Talamancana*, to include *P. lonera*^41^. A second, large and apparently well-defined clade contains *P. zerlina* and relatives, all of which have a very distinctive synapomorphy, a keel-like spine on the dorsal side of the aedeagus near the posterior tip (character 1:1). Genitalia barely differ among any of the eleven species in this clade, although the adult mimetic wing patterns and the immature stage morphology and biology show striking differences.

### New calibration of the Solanaceae phylogeny

To calibrate the phylogeny of *Pteronymia*, we used a combination of secondary calibrations of Nymphalidae ages^38^ and maximum age constraints based on host-plant lineage ages (Solanaceae). To extract those ages, we used the molecular matrix of a previous Solanaceae phylogeny^42^ to generate a new phylogeny, which was calibrated with the stem age of the family extracted from a dated phylogeny of Angiosperms^43^. Our Solanaceae phylogeny shows similar node support and topology to those of the latest published phylogeny of Solanaceae^42^. Median lineage ages are on average about 25% older and have wider 95% credibility intervals (Table 2, see Supplementary Fig. S4 online). In most lineages the 95% credibility intervals presented here span the median age of the published phylogeny of Solanaceae^42^, but the median ages themselves fall almost always outside the 95% credibility intervals of the previous phylogeny (Table 2). The wider credibility intervals in our study are due to the use of the more conservative uniform prior on the calibration point as compared to the previous phylogeny of Solanaceae, which implemented a log-normal prior that tends to drive ages towards the mode of the prior distribution^42^. Consequently, the ages of the lineages used for calibrating the phylogeny of *Pteronymia* are older than those inferred previously^42^ (Table 2).

**Table 2.** Comparison of our estimates of ages of Solanaceae lineages with those of Särkinen et al. (2013)42. *ages used as calibration in the *Pteronymia* phylogeny.

| Solanaceae clades | Age estimates<br>This study | Age estimates<br>Särkinen et al. (2013) |
| --- | --- | --- |
| Stem of Solanaceae* | 64.0 my [48 – 83] | 49my [46 -54] |
| Crown of Solanaceae | 42.2 my [26.8-55.2] | 30.3 my [26.3-34.0] |
| Crown of “x=12” clade (MRCA *Solanum* and *Nicotiana*) | 39.5 my [21.9-44.4] | 23.7 my [23.0-25.7] |
| Crown of *Solanum* | 20.9 my [14.5-29.5] | 15.6 my [13.1-17.5] |
| Stem of *Capsicum* * | 19.0 my [11.5 - 25] | 13 my [10 – 16] |
| stem of *Schultesianthus* * | 17.0 my [7 – 24] | 13 my [9 – 19] |
| stem of *Brugmansia* * | 10.5 my [5 – 14] | 7 my [4 – 10] |

### *Dated combined* Pteronymia *phylogeny*

We combined morphological and molecular data to generate a species-level phylogeny that was calibrated using secondary calibrations from a published Nymphalidae phylogeny^38^, and from Solanaceae lineage ages estimated in this study, using BEAST 1.7.5^44^. The combination of morphological and molecular data generated a phylogeny comprising all known extant species (Fig. 1). Many nodes were poorly supported, and this was mostly caused by the species represented only by morphological characters, whose placement was uncertain. Our calibration strategy that combined butterfly- and host-plant-derived secondary calibrations resulted in ages that were about 30 to 50% older than those inferred in a recent time-calibrated phylogeny of Ithomiini genera that relied on previous minimum age estimate of Solanaceae lineages^37^. By contrast, our ages were often slightly younger, but well within the credibility interval of the ages estimated in the Nymphalidae phylogeny^38^. Notably, the stem age of *Pteronymia* inferred in our study is 14.4 million years (my) [12.3 – 16.3], while it was 7.5 my [6.0 – 9.0] in the higher level Ithomiini phylogeny^37^ and 15.7 my [11.5 – 18.5] in the Nymphalidae phylogeny^38^. We repeated this analysis without the Solanaceae calibrations and this had little impact on most nodes of the phylogeny. The greatest difference was found for the divergence between the outgroup genera *Athesis* and *Patricia* (22.9 million years ago (mya) [20.0 – 24.0] under the combined calibration scheme, 25.4 mya [21.8 – 27.5] with Nymphalidae calibrations only). For the genus *Pteronymia*, ages under the two calibration schemes were extremely similar (regression ages Nymphalidae calibration only (N) versus combined Nymphalidae and Solanaceae (S): N = 0.989*S, r2=0.997). Trees generated under the combined calibration scheme were used in all subsequent analyses.

**Figure 1:**
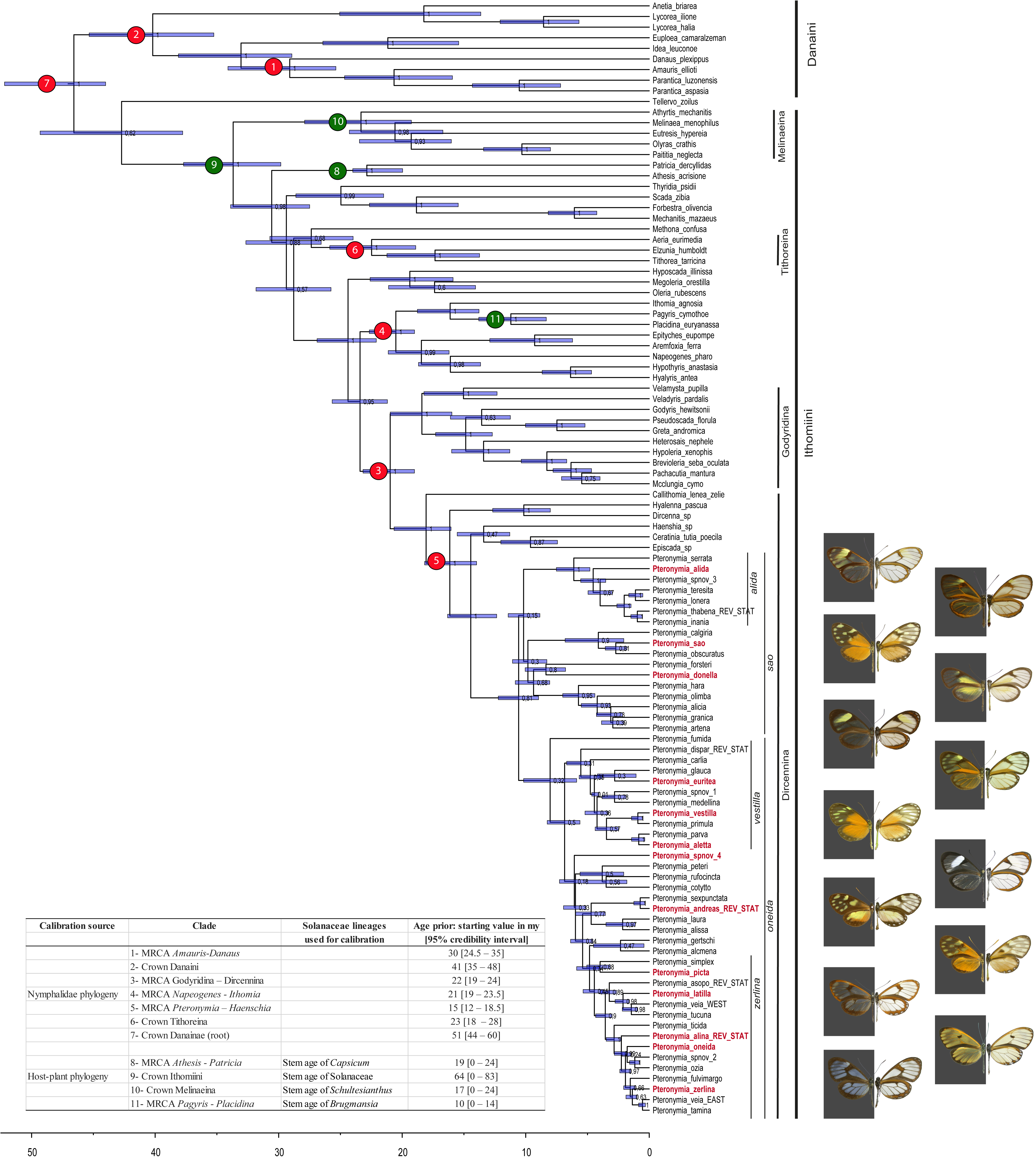
BEAST dated species-level phylogeny of the genus *Pteronymia*, based on molecular and morphological characters. Main clades and secondary calibration points based on butterfly (red circles) and host-plant ages (green circles) are indicated and corresponding age priors are shown in the table inserted in the figure. The figure was generated with FigTree (http://tree.bio.ed.ac.uk/software/figtree/) and edited with Adobe Illustrator 4 (http://www.adobe.com/uk/products/illustrator.html). Butterfly pictures were taken by Keith Willmott and edited in Adobe Photoshop CS4 (www.adobe.com/products/photoshop/).

After splitting from its sister lineage (the clade composed of the genera *Episcada*, *Ceratinia*, and *Haenschia*) 14.4 mya [12.3 – 16.3], the genus *Pteronymia* started diversifying about 10.6 mya [9.0 –12.2], when it split into two major, but poorly supported, clades: the *P. sao* clade, 17 species, and the *P. oneida* clade, 36 species. The two main clades appearing in the morphological phylogeny were also recovered in the combined analysis, with the exception that *P. latilla* and *P. tucuna* grouped with two species in the *P. zerlina* clade. The male genitalia of both of the former species are rather different to remaining members of the *P. zerlina* clade, lacking the distinctive male genitalic synapomorphy of that clade in addition to also lacking a gnathos, the loss of which occurs within the genus only in *P. latilla*, *P. tucuna*, *P. sao* and *P. obscuratus*. Otherwise, none of the relationships implied in the combined phylogeny seem to contradict any strong morphological evidence.

Because of topological uncertainty of the *Pteronymia* phylogeny, we performed subsequent analyses both on the MCC tree and on a random subset of 100 trees from the posterior distribution.

### Spatial patterns of diversification

We used georeferenced records (Supplementary Fig. S5) to analyse the spatial patterns of diversification of *Pteronymia* across nine biogeographic areas (Fig. 2). We performed biogeographical analyses using the software RASP 2.1^45^. The analyses on the MCC tree and on the 100 trees yielded very similar results (Fig. 3, see Supplementary Fig S6 online), and only the analyses on the MCC tree are presented here. Our biogeographic reconstruction suggests that the most likely ancestral area for the genus *Pteronymia* is the Western/Central Northern Andes (hereafter, Northern Andes), i.e., the area comprising the slopes of the Western and Central cordillera of Colombian (and Ecuadorian) Andes (Fig. 3), although there is uncertainty as to whether the origin of the genus was limited to this area, or also spanned neighbouring regions (see Supplementary Fig S7 online). The two main clades (*P. oneida* and *P. sao* clades) also originated and started diversifying in the same area, with some uncertainty as to whether ancestral lineages spanned larger regions for the *P. oneida* clade (see Supplementary Fig. S6 online). A large proportion (55%) of speciation events occurred within the Northern Andes. In particular, the most likely ancestral area for the young and diverse *P. zerlina* clade was the slopes of the Northern Andes. Rapid diversification subsequently occurred within the last 3.6 my [3.0 – 4.3] in this clade, which is coincident with the final uplift of the Eastern Cordillera of Colombia and the Venezuelan Cordilleras c. 5-2 mya^9^. The Central Andes appear relatively species poor compared to the Northern Andes; only 13 species occur in the Central Andes. These are the result of multiple independent colonization events (10 events), and to a much lesser extent from local diversification (e. g., the clade encompassing *P. hara* and its sister clade, three species in this region). The oldest colonizations of the Central Andes were recovered in the *P. sao* clade (in the last 5.8 my [4.4 – 7.0]), where such colonizations happened at least five times.

**Figure 2:**
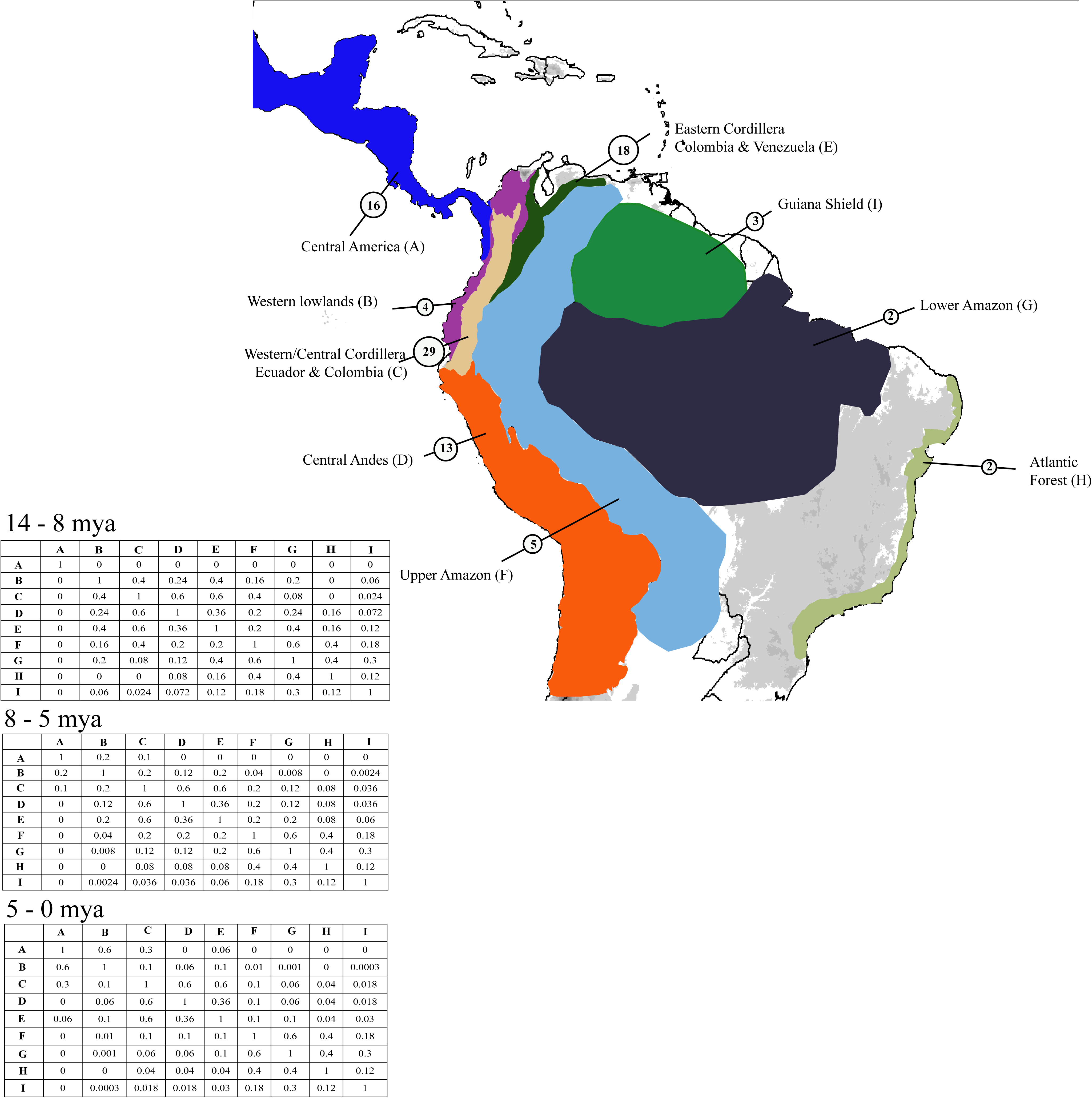
Biogeographical regions used in the DEC and SDEC models for reconstruction of ancestral areas and the dispersal probability matrices for the different time slices. Species richness in each areas are shown in the circles. The map was generated using ArcGIS 9.3 (http://www.edit.com/software/arcgis/), and edited with Adobe Illustrator 4 (http://www.adobe.com/uk/products/illustrator.html).

**Figure 3:**
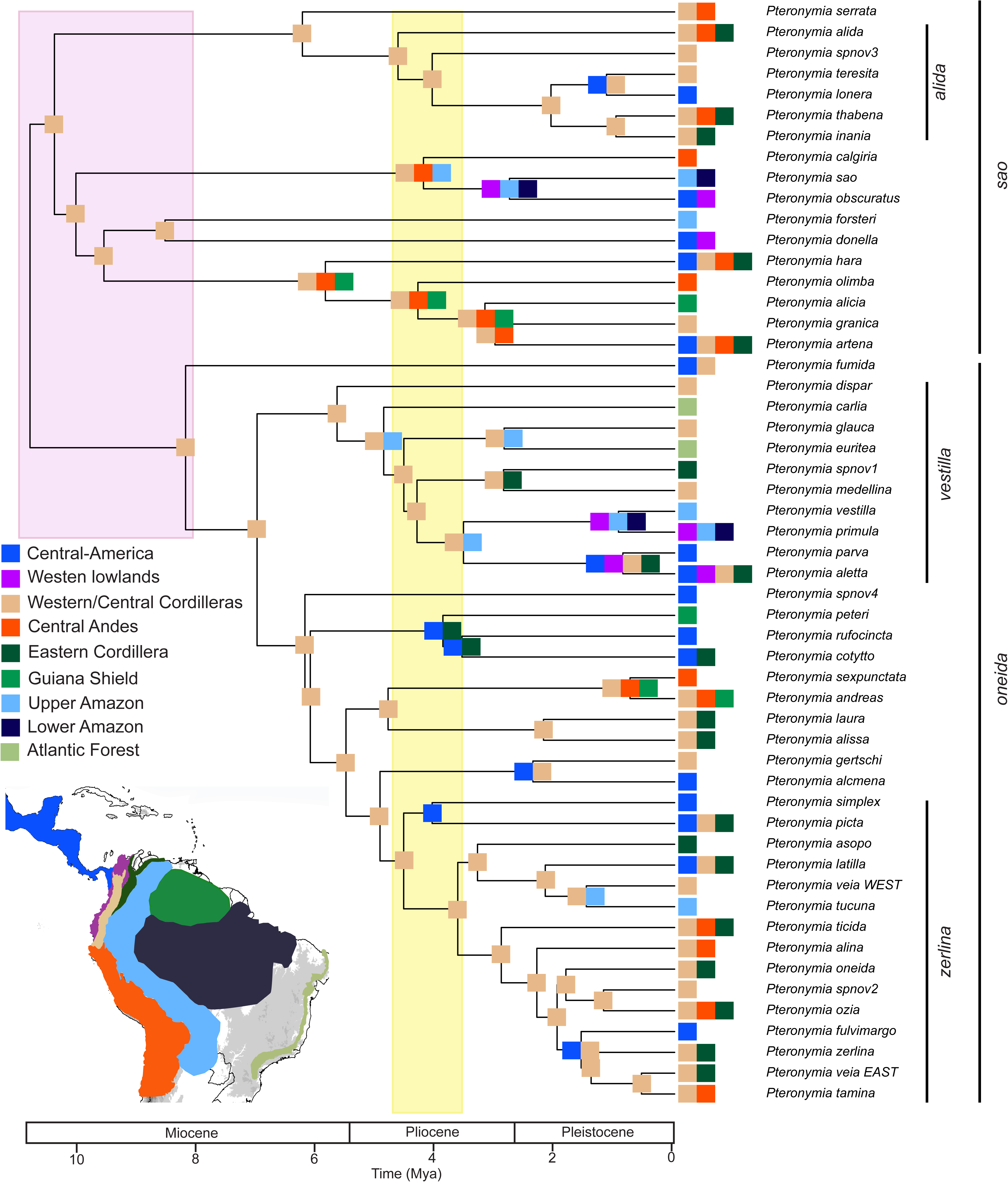
RASP historical biogeography inference (best maximum likelihood estimates on the MCC tree). Major paleoenvironmental events are indicated by large coloured rectangle (light pink: drainage of te Pebas system; light yellow: hypothesized closure of the Isthmus of Panama). The figure was generated with R (https://cran.r-project.org/) and edited with Adobe Illustrator 4 (http://www.adobe.com/uk/products/illustrator.html).

A small number of lineages colonized the Upper and Lower Amazon regions. Only one dispersal event was followed by local diversification in Amazonia, in the *vestilla* clade (the clade encompassing *P. dispar* and its sister clade). Three other colonizations of Amazonia occurred independently throughout the phylogeny, resulting in an overall very low Amazonian diversity, particularly in the lower Amazon. All these events happened within the last 5.6 mya [4.8 – 6.7]. The two species that colonized the Atlantic Forest arose from Amazonian lineages, and did not result in local diversification. Conversely, a high number of independent colonization events (15, according to the maximum likelihood estimates) occurred from the Andes to Central America. Most of those colonizations occurred within the last 5.0 my [4.0 – 5.9]. Colonization time of Central America was highly uncertain for *P. fumida* and *P*. spnov 4, because these species split from their sister lineages 10.6 mya [9.0 – 12.2] and 7.0 mya [5.6 – 8.3], respectively, and may have therefore colonized Central America any time during those periods. Colonization of Central America was sometimes, but rarely given the high number of colonizations, followed by local speciation (e.g. *P. lonera* and *P. teresita*, *P. alcmena* and *P. gertschi*).

### Evolution of elevational range

We investigated the evolution of the elevational range and mean elevation of *Pteronymia* species on the MCC tree and on 100 trees from the posterior distribution. The phylogenetic signal and the tempo of evolution of the elevational range and mean elevation were assessed by estimating simultaneously the maximum likelihood values of the λ and δ branch scaling parameters^46^, respectively. A λ value of one indicates that the phylogeny correctly represents the trait covariance among species (Brownian motion model of evolution), while λ < 1 indicates that the phylogeny overestimates the trait covariance among species. A δ value of one means that the trait evolves at a constant pace along branches of the tree; δ < 1 indicates early changes in the character values followed by a slowing down of the evolution rate; while δ > 1 indicates accelerated evolution rate and species-specific adaptation. For the mean elevation and the elevational range, estimates of λ across all trees from the posterior distribution and for the MCC tree were not significantly different from one, meaning that trait evolution does not differ from a Brownian motion model. Estimates of δ for the mean elevation were also not significantly different from one (Table 3). For the lower and upper boundaries of the elevational range, the estimates of δ were significantly higher than one in >50% of the trees of the posterior distribution (Table 3). The MCC tree showed significantly higher estimates of δ for the lower boundary of the elevational range, but only marginally significant for the upper boundary (Table 3). Values of δ higher than one indicate an acceleration of the rate of evolution of elevation range and species elevational specialisation. Results differ between the mean elevation and elevational range perhaps because the mean is less variable than the range (e.g., species with different elevational ranges may have similar elevation mean). Reconstructions of ancestral mean elevation and range boundaries accounting for the inferred δ values are depicted on Fig. 4.

**Figure 4:**
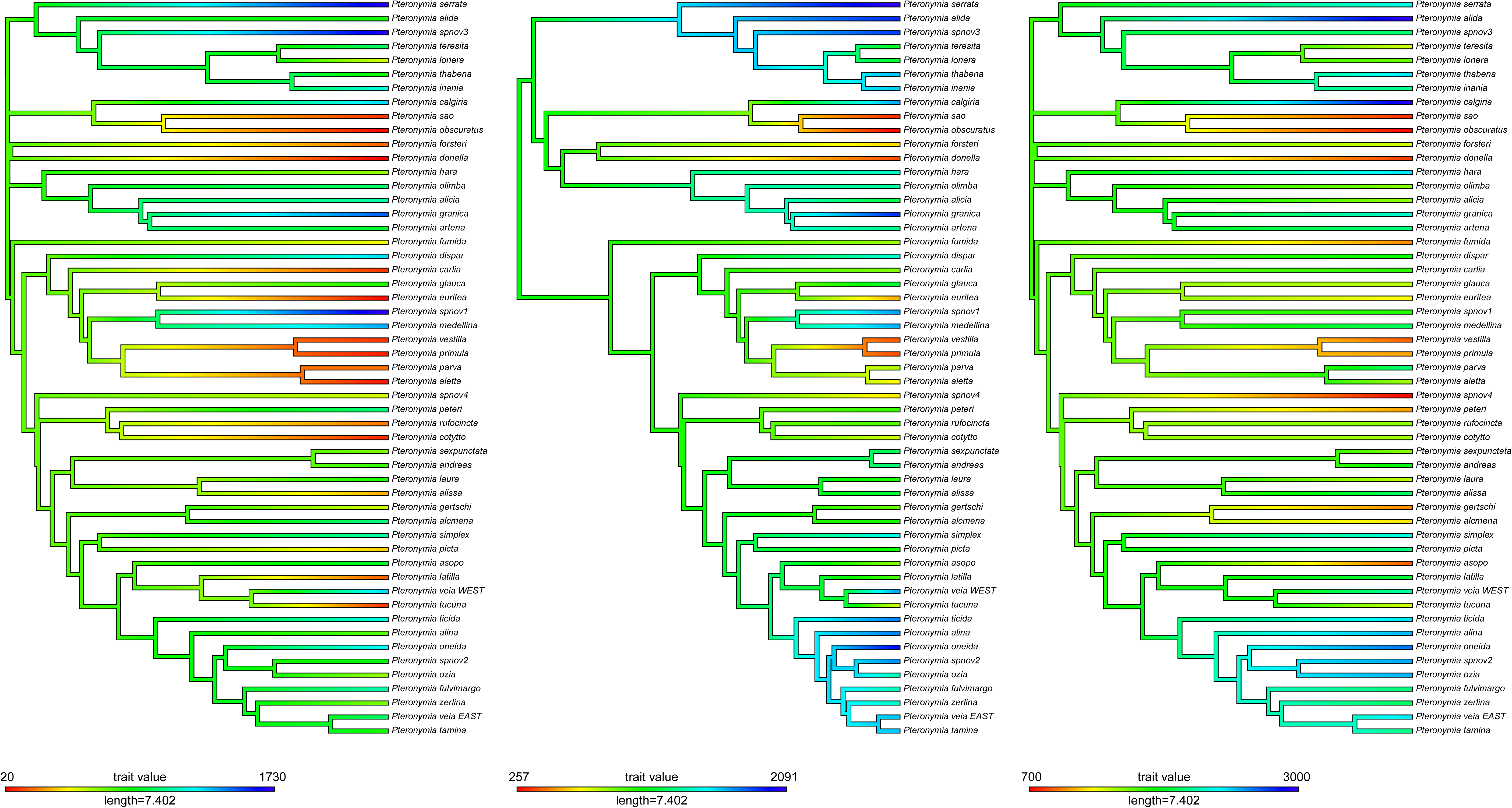
Ancestral reconstruction of the mean and boundaries of the 95% elevational range (left: lower boundary, middle: upper boundary, right: mean). For the lower and upper boundaries of the elevational range, trees were rescaled according to the δ value inferred (Table 2). The figure was generated with R (https://cran.r-project.org/).

**Table 3.**
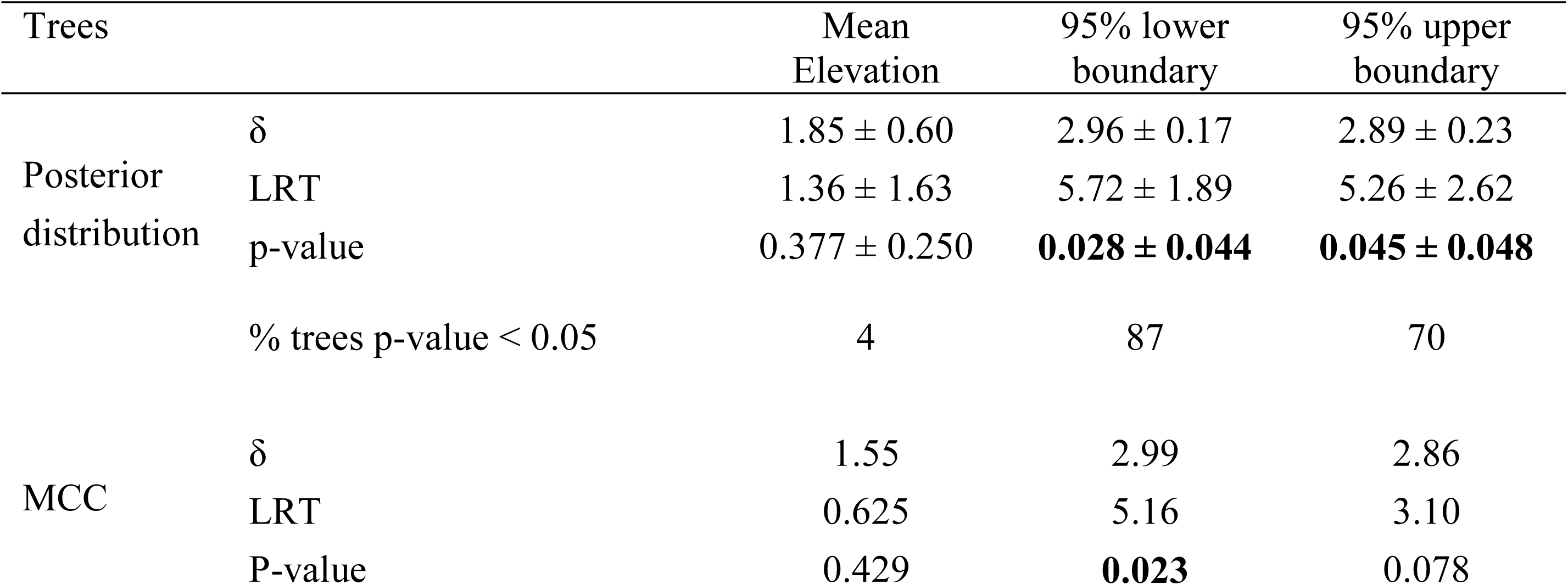
Maximum likelihood estimates of δ for the mean and boundaries of the 95% elevational range, for the 100-tree posterior distribution (average values ± standard deviation), and for the MCC tree. Likelihood Ratio Tests values when compared with the null model (δ = 1), corresponding p-values and fraction of trees for which δ is significantly different from 1 are reported.

### Temporal patterns of diversification

We first investigated heterogeneity among clades for speciation and extinction rates using MEDUSA^47
^ on the MCC tree and on 100 trees from the posterior distribution. No significant shift of diversification rates was found on the MCC tree. Five trees of the posterior distribution (5%) had at least one significant shift of diversification rates. Each of those shifts was found in less than 5% of the trees, and therefore considered as non-significant. We then investigated whether diversification rates varied through time by fitting time-dependent models of speciation and extinction rates^48^. The best fitting model was an exponential time-dependent speciation rate without extinction (Table 4). According to this model speciation rate decreased from 0.646 event per lineage per million year for the MCC tree and 0.538 ± 0.024 for the 100 trees at the origin of the genus (crown) to 0.148 for the MCC tree and 0.159 ± 0.002 for the 100 trees at present (Fig. 5). All other models, including the constant speciation rate model, were rejected at the threshold of ΔAIC > 2, strengthening the support for a decreasing speciation rate through time.

**Table 4.** Models of time-dependent diversification fitted on 100 trees from the posterior distribution, ranked by increasing AIC score. Mean values of parameters are indicated followed by the standard deviation in brackets. BCST=constant speciation, BVAR=time-dependent speciation, DCST=constant extinction, DVAR time-dependent extinction. logL = likelihood of the model, AIC = AIC score, ΔAIC = difference of AIC between the each model and the best fitting model, λ=speciation rate at present, α=coefficient of time variation of the speciation rate, μ=extinction rate at present, ß=coefficient of time variation of the extinction rate.

| Model | Par | logL | AIC | ΔAIC | $\lambda$ | $\alpha$ | $\mu$ | $\beta$ |
| --- | --- | --- | --- | --- | --- | --- | --- | --- |
| **BVAR** | **2** | **-122.93**<br>**(0.286)** | **249.86**<br>**(0.571)** | **0.00** | **0.161**<br>**(0.001)** | **0.110**<br>**(0.003)** |  |  |
| BVAR - DCST | 3 | -122.93<br>(0.286) | 251.86<br>(0.571) | 2.00 | 0.161<br>(0.001) | 0.110<br>(0.003) | 5.00E-08<br>(5.00E-08) |  |
| BCST | 1 | -125.39<br>(0.258) | 252.79<br>(0.516) | 2.93 | 0.233<br>(0.001) |  |  |  |
| BVAR - DVAR | 4 | -122.93<br>(0.286) | 253.86<br>(0.571) | 4.00 | 0.161<br>(0.001) | 0.110<br>(0.003) | 5.00E-08<br>(5.00E-08) | -0.004<br>(0.003) |
| BCST - DCST | 2 | -125.39<br>(0.258) | 254.79<br>(0.516) | 4.93 | 0.233<br>(0.001) |  | <1.00E-08<br>(<1.00E-08) |  |
| BCST - DVAR | 3 | -125.39<br>(0.258) | 256.79<br>(0.516) | 6.93 | 0.233<br>(0.001) |  | <1.00E-08<br>(<1.00E-08) | 0.005<br>(2.00E-04) |

**Figure 5.**
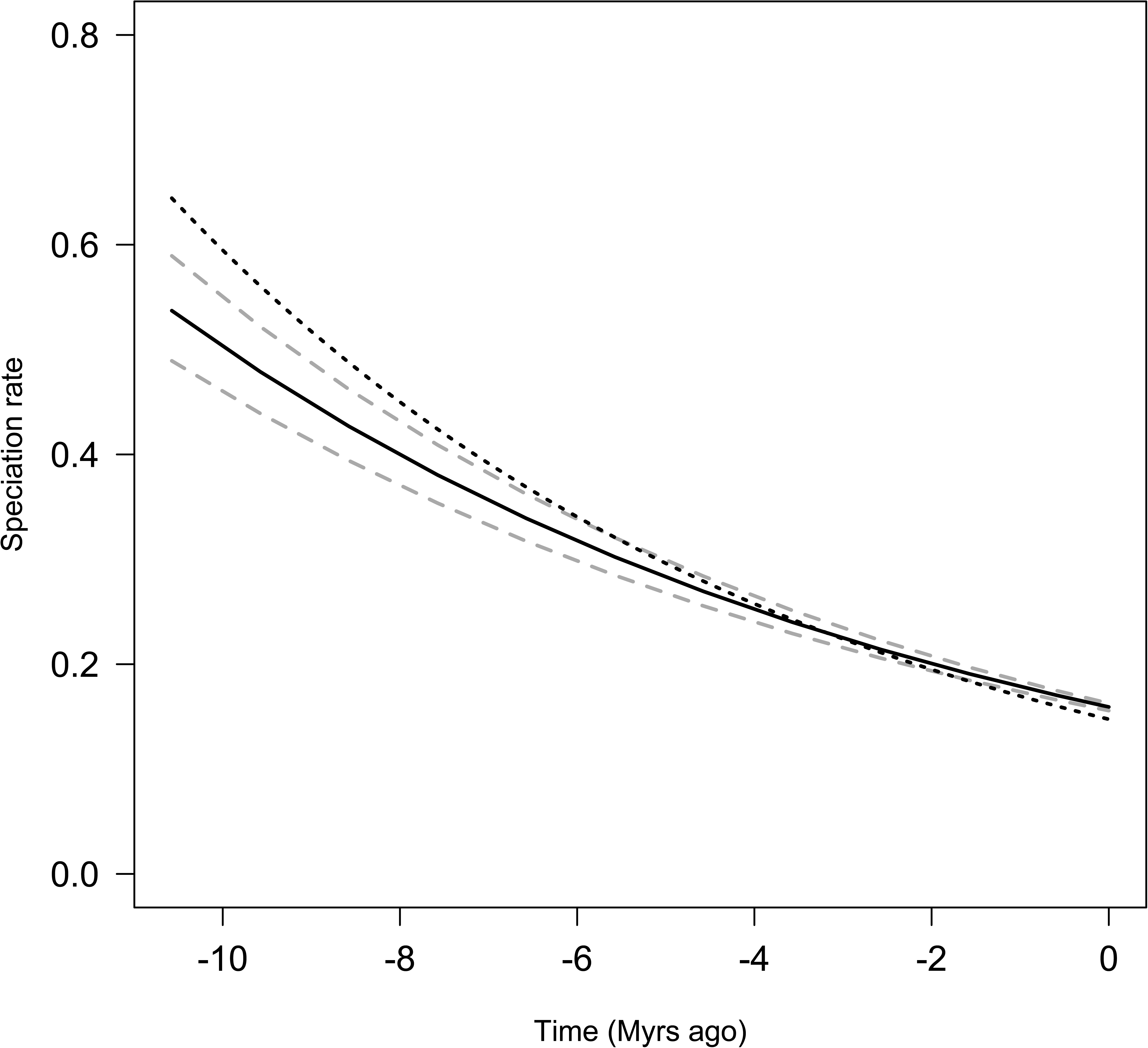
Speciation rate through time estimated by the best fitting model of diversification (Table 4). The model was fitted on the MCC tree and on 100 trees. The dotted line corresponds to the speciation rate of the MCC tree. The plain line corresponds to the mean speciation rate according of the 100 trees, and dashed lines correspond to the 95% confidence interval. The figure was generated with R (https://cran.r-project.org/).

## Discussion

Our extensive molecular sampling of the genus *Pteronymia* encompassing multiple subspecies and combined with morphological and distributional data enabled us to redefine species boundaries in the genus, resulting in six additional recognized species in the genus. The resulting taxonomic changes now make *Pteronymia* the most species-rich ithomiine genus, with 53 species. Morphological characters that are typically useful in diagnosing ithomiine species, such as in the genitalia and wing androconia, were clearly unhelpful in these cases, although mimetic wing pattern and life histories, where known, often did show significant variation. As knowledge of the biology of different populations and more material for molecular study become available, additional changes to the species taxonomy may be needed in future. At deeper levels of the phylogeny, despite *Pteronymia* showing some of the greatest diversity within the Ithomiini in terms of the male and female genitalia, wing venation and androconia, immature stage morphology and biology, this variation proved remarkably unhelpful in resolving the phylogeny, perhaps due to high rates of morphological character evolution. The combination of morphological and molecular characters obviously increased the resolution of the phylogeny compared to morphology alone, but many clades were still surprisingly poorly supported, and in any case the phylogeny of *Pteronymia* was less resolved than those of other ithomiine genera^6,27,49,50^, even when considering only molecular characters. This may be due to incomplete lineage sorting, rapid diversification or hybridization. While hybridization and introgression seem common in other mimetic butterflies, such as *Heliconius*^51–53^, nothing is known about such processes in ithomiine butterflies. Future genomic data may shed light on demographic processes and gene flow between species, which may have contrubted to the poor support seen in some nodes in the *Pteronymia* phylogeny.

Our calibration strategy, based on a combination of secondary calibration derived from host-plant and Nymphalidae phylogenies, required the recalibration of a published Solanaceae phylogeny^42^ (host-plants of most Ithomiini) because original ages were biased toward present, which precludes using those data as maximum calibrations. Our new calibration scheme, based on a secondary calibration extracted from a fossil-dated phylogeny of Angiosperms^43^, inferred ages of Solanaceae lineages that were about 25% older than previous estimates^42^, but mostly within the 95% confidence range of those estimates. The host-plant ages used in this study as maximum constraints are based on one of the ‘youngest’ hypothesis for Angiosperm diversification and recent studies suggest older ages of Solanaceae. A phylogenetic inference using genomic data found a much older origin of Angiosperms^54^, but this had a moderate impact on the stem age of the order containing Solanaceae (Solanales: ca. 92 mya^54^ versus 85.9 mya^43^, with overlapping 95% confidence ranges^54^) and presumably on that of Solanaceae (not inferred in that study). The recent description of a 52.2 my old *Physalis* fossil^55^, a solanaceous genus that is inferred to be 9.1 [5.9 – 12.9] my old in our study (stem age), is much more challenging for the ages of Solanaceae and Angiosperms as a whole. It is possible that this recently described *Physalis* fossil represents an earlier lineage of Solanaceae. The inflated calyx is found in many genera throughout the ‘berry’ clade of Solanaceae (i. e., the subfamily Solanoideae, where the stem is the MRCA of *Nicotiana* and *Solanum* and the crown the MRCA of *Latua* and *Solanum*^56^) and transcription factors governing this character appear to be plesiomorphic in the family^57–59^. Placement of the fossil at an earlier diverging node, such as that of the berry clade, within Solanaceae is less contradictory in terms of Angiosperm evolution as a whole, and would not have a major effect in pushing back the stem and crown node age of Solanaceae. In terms of our findings here, older Solanaceae age estimates does not affect our time-calibrated *Pteronymia* tree given that butterfly clades appear to be much younger than their corresponding host-plant groups even when host-plant ages were inferred from one of .the ‘youngest’ Angiosperm time-calibrated phylogenies^43^.

Our combined calibration strategy therefore resulted in ages that were consistent with those inferred in the Nymphalidae phylogeny^38^, but older than those inferred in the higher-level Ithomiini phylogeny^37^. This difference stems from several factors. The new ages of the Solanaceae lineages (this paper) used as maximum calibration were 25% older than the previous estimates used in the higher-level Ithomiini phylogeny^37^, and we used the oldest boundary of the 95% credibility interval as the (hard) maximum age of corresponding ithomiine lineages, instead of the mean as in the higher-level Ithomiini phylogeny^37^. The rationale for this is that minimum (such as fossil-based) and maximum (such as host-plant-based) calibrations ought to be conservative and therefore account for uncertainty in calibration age^60^. In our case, where we implemented conservative uniform priors between maximum ages and present, node ages in the *Pteronymia* phylogeny were not necessarily attracted towards maximum ages, they were just allowed to go as far as those ages. Since host-plant ages provide maximum calibrations, they need to be combined with minimum calibrations, in a way similar to fossil-based minimum calibrations that need to be combined with at least one maximum calibration point^60^. Given the absence of fossils for Danainae, here we supplemented the host-plant-derived calibrations with secondary calibrations extracted from the Nymphalidae phylogeny^38^, which was calibrated with a combination of fossils and host-plant constraints. These calibrations provided both minimum and maximum ages, and therefore imposed a stronger prior on ithomiine ages than the host-plant-derived calibrations, but to be as conservative as possible we used a uniform prior spanning the 95% credibility interval of the Nymphalidae ages that were used for calibration.

Our results based on the biogeographic reconstruction of the genus *Pteronymia* and evolution of elevational range were consistent across trees of the posterior distribution despite the relatively poor resolution of the MCC phylogeny, and revealed the fundamental roles played by the Northern Andes in the diversification of the group, in multiple ways. The Northern Andes have probably been (1) the area of origin of the group (although the inference for the root is not well resolved, this region appears in all the potential ancestral areas); (2) the centre of early and sustained local diversification; and (3) a source of recent colonizations to lowland areas and to Central America, that led to an accelerated evolution of the elevational range.

The biogeographical pattern of diversification, where the Andes play a central role, resembles those found in other Andean ithomiine genera, such as *Hypomenitis*^34^, and to a lesser extent *Napeogenes*^6^ and a clade of *Oleria*^4^. These groups originated and diversified in the Northern Andes during the last 10 million years, a period during which the Northern Andes experienced different phases of intense uplift^9^. This period of orogeny involved great landscape transformations that may have affected the dynamics of Andean lineages, by isolating populations within deep valleys or on both sides of the Andes, but also in modifying the climatic conditions. Also, the slopes of the Andes offer a great number of opportunities for ecological speciation due to significant environmental turnover resulting in high habitat, host-plant and predator diversity. In the genus *Pteronymia*, the observed decrease of speciation rate with time and the low support for basal nodes may indicate a rapid diversification driven by adaptive factors. Elevations up to 2000m probably already existed by 10 million years ago in the Northern Andes^61^. Given the timing of diversification of the genera *Hypomenitis, Napeogenes* and *Pteronymia* in the last 15 to 5 million years, the majority of speciation events have likely not coincided with the appearance of newly available elevations. Instead, speciation was probably facilitated by an already well-established and diverse ecosystem. The slopes of the Andes harbour not only a diversity of habitats and host-plant communities^62,63^, but also a diversity of mimicry rings^27^. *Pteronymia* is one of the most diverse ithomiine genera in terms of mimetic wing colour patterns. Shifts in colour pattern are known to drive speciation in other mimetic butterflies such as *Heliconius*^*64,65*^, where colour pattern is considered as a ‘magic’ trait, i.e., a trait that is both under disruptive selection and associated with assortative mating^64,66,67^. Mimetic butterflies often harbour multiple geographic races with different colour patterns, and it has been shown in *Heliconius* butterflies that interracial hybrids that display an intermediate, non-mimetic colour pattern suffer higher predation^65^. Shifts in colour pattern may therefore cause postzygotic reproductive isolation. In addition, *Heliconius* butterflies tend to prefer mates with their own colour pattern over conspecifics with a different colour pattern^64,66,67^, thereby driving prezygotic reproductive isolation. In this case, loci involved in mate preference and in colour pattern are tightly linked^68^. In Ithomiini, experimental evidence for the role of colour pattern as a mating cue is absent due to the difficulty of maintaining and rearing Ithomiini in captivity, but observation suggests that this may be the case^69^. Moreover, shifts in colour patterns have been shown to be statistically associated with cladogenesis in phylogenies^35,70^, consistent with a role of colour pattern in reproductive isolation. Shifts in colour pattern may therefore have contributed to Andean diversification in the genus *Pteronymia*.

In contrast with other ithomiine clades^4,6,34^, little local diversification in *Pteronymia* occurred outside of the Northern Andes. An exception to this is the *P. vestilla* clade, which expanded into and partly diversified in lowland areas. According to our reconstructions, this lineage dispersed into the Upper Amazon from the Northern Andes and started diversifying locally around 5.6 mya [4.8 – 6.7]. Local diversification was then followed by expansions or dispersal into other areas, including the lower Amazon and Guiana shield, the Atlantic forest, and Western lowlands of Colombia and Ecuador. Only two extant lineages of this clade occur in Amazonia. The Upper Amazon region has experienced important environmental changes since the Miocene. During most of the Miocene, the region was covered by the large lake and shallow swamps of the Pebas system, which was connected northward to the Caribbean Sea and potentially westward to the Pacific Ocean^9^. This particular ecosystem may have had a major influence on the diversification of clades restricted to forest habitats^14,22^ by limiting occurrence and therefore speciation to the margins of this region, but probably also by preventing dispersal between Amazonia and the Andes and between the Northern and the Central Andes (via the West Andean Portal, a low elevation gap between the Northern and Central Andes^71^). The demise of the Pebas system started around 10-8 mya and it was rapidly drained eastward, leading to the establishment of the modern Amazon basin, probably around 7 mya^9^. Our results suggest that dispersal and diversification in Amazonia did not occur simultaneously with the drainage of Pebas, but later on. The diversification of the *P. vestilla* clade in the Upper Amazon occurred fairly recently (5.6 mya [4.8 – 6.7]), as did the independent colonizations of the Upper Amazon by the ancestor of *P. veia_*WEST and *P. tucuna* (1.5 mya [0.9 -2.2]), and the ancestor of *P. sao* and *P. obscuratus* (2.7 mya [1.8 – 3.6]). The timing of the colonization of the Upper Amazon by *P. forsteri*, which split from its sister clade 8.3 mya [6.8 -10], is much less precise, since it could have happened any time during this period.

Because little local diversification in *Pteronymia* occurred outside of the Northern Andes, most of the diversity in non-Andean regions is due to colonization out of the Andes. Species diversity in the Central Andes and Central America built up through the accumulation of independent dispersal events, rarely followed by speciation events. Many of these events occurred without strong elevational shifts suggesting that dispersal may have been facilitated by the existence of similar ecological conditions in montane areas. By contrast, dispersal toward lowland areas, such as the Upper Amazon, may have entailed more adaptations to fit different bioclimatic conditions or host-plants, which likely explains the rare occurrence of such events.

There is a debate surrounding the timing of the closure of the Panama Isthmus. The hypothesis that it occurred very recently (5-3 mya) has been widely adopted in the literature (see^12^. However, both geological and paleontological findings suggest a possible much earlier appearance of land masses, possibly as early as the early or middle Miocene (e.g.^12,72–74^. In our biogeographical analysis, although interchanges between Central America and South America were allowed earlier, most of the colonization events of *Pteronymia* lineages toward Central America have occurred during the last 5.0 [4.0 – 5.9] million years, in agreement with the first hypothesis. However, the colonization time of Central America by *P. fumida* and *P.* spnov 4, two species with long branches, is uncertain, and may have happened much earlier.

The pattern of diversification inferred for *Pteronymia* is very similar to biogeographic patterns of other taxa described in the literature. For example, in vertebrates, the Northern Andes were a major centre of diversification for glassfrogs (Allocentroleniae), which subsequently fed the adjacent areas through dispersal, including the Central Andes and the non-Andean regions^3^. A similar conclusion was reached in the Thraupini tanagers, which also diversified during the last 10 million years, with higher rates of colonization out of the Northern Andes (mostly toward the Central-Andes) rather than into that region^75
^. The bat genus *Sturnina* also diversified during the last 10 million years, from the Northern Andes toward the rest of the Neotropical region^76^. Similarly, in plants, many studies report the Northern Andes as a centre of diversification and a source for adjacent areas^23^. The Rubiaceae subfamily Cinchonoideae originated and mostly diversified in the Northern Andes, with subsequent colonization of both lowland and highland adjacent areas^71^. Extremely high rates of diversification were reported in the Andes for the genus *Lupinus*^7^. The Campanulaceae experienced higher rates of diversification in the Andes that were correlated with paleoelevations of the Andes^8
^, suggesting that the Andes have directly affected diversification in this family. Taken together, these findings are consistent with the idea that, at least during the last 10 million years, the Northern Andes have been an important source of biodiversity in the Neotropics probably due to geological, climatic and biotic factors, with local diversification followed by lineage dispersal throughout other Neotropical regions.

## Methods

### Morphological characters and phylogeny

Prior to our study, 47 species were listed in the genus *Pteronymia*, but this figure increased to 53 after taxonomic revision based on new data (Table 1). Forty-six adult and 41 immature (i.e., larval and pupal) morphological characters were examined in 52 *Pteronymia* species (after our revision, see Supplementary Methods S1, Supplementary Table S2, Supplementary Figs. S8-S9 online; *P. dispar***rev. stat.** was not coded) and two outgroup species (*Episcada apuleia* and *Dircennina adina*).

A Maximum Parsimony topology was estimated in TNT v. 1.5-beta^77^ using the New Technology Search, with all four search methods – ratchet, tree-fusing, tree-drifting and sectorial, with default parameters, 100 random additional sequences and random seed equal 0, with all characters equally weighted. The majority-rule consensus tree, consistency index (CI) and the retention index (RI) were computed in Winclada^78^. The stability of each branch was estimated using the non-parametric bootstrap test, with 1,000 replicates and 100 random taxon additions, and Bremer support, using the script Bremer.run in TNT.

### Molecular characters and phylogeny

We used a total of 166 *Pteronymia* specimens for molecular analyses, representing 41 of the species recognized prior to our revision, and 47 of the species recognized after our revision (Table 1; see Supplementary Table S1 online). Species with no molecular data were *P. alcmena, P. alicia, P. calgiria,P. fumida, P. glauca* and *P. peteri*.

We used *de novo* (ca. 85%) and published (ca. 15%) sequences from five gene regions to infer a molecular phylogeny (see Supplementary Table S1 online): the mitochondrial region spanning the mitochrondrial genes *cytochrome oxidase c subunit 1*, *leucine transfer RNA* and *cytochrome oxidase c subunit 2* (CO1, tRNAleu, CO2, 2356 bp), and the nuclear genes *tektin* (735 bp) and *Elongation Factor 1 alpha* (EF1A, 1259 bp). Species coverage was 98% for the mitochondrial fragment, 73% for *tektin* and 63% for EF1A. Primers and PCR conditions followed previously described conditions^6^. In addition, 52 ithomiine and danaine outgroup species were selected^34^ (see Supplementary Table S1 online). The dataset was then partitioned by gene and codon positions and the best models of substitution for optimized sets of nucleotides were selected over all models implemented in (1) RAxML^79^ and (2) MrBayes^80^, using the ‘greedy’ algorithm and linked rates implemented in PartitionFinder 1.1.1^81^ (see Supplementary Table S3 online).

We performed a maximum likelihood phylogenetic inference using RAxML^79^ on the Cipres server^82^. In addition, we performed a Bayesian inference of the phylogeny using MrBayes 3.2.2^80^ on the Cipres server^82^. Substitution models of each partition were re-estimated in MrBayes 3.2.2 using the reversible-jump MCMC^83^. Two independent analyses were run for 10 million generations, with four Monte Carlo Markov chains each and a sampling frequency of one out of 10,000 generations (resulting in 1,000 posterior trees). After checking for convergence, the posterior distributions of the two runs were combined, with a burnin of 10%. The maximum clade credibility tree with median node ages was computed using TreeAnnotator 1.6.2^44^. The resulting tree was used to investigate topology and species boundaries.

### Molecular dating

In order to estimate a time-calibrated phylogeny of *Pteronymia*, we combined two types of time constraints: the age of larval host-plants and age estimates from higher-level phylogenies of butterflies. Host-plant ages can be used as maximum age constraints in phylogenies of mono- or oligophagous herbivores^37,38^, assuming that such herbivorous taxa diversified only after the emergence of their host-plant lineages. Most Ithomiini feed on Solanaceae, which represents a host-plant shift from the ancestral host-plants of Danainae (Apocynaceae^84^). In a previous study aiming at dating a higher-level phylogeny of the tribe Ithomiini^37^, ages of several Solanaceae lineages inferred from a dated phylogeny of Solanaceae^42^ were used as maximum calibrations for Ithomiini clades feeding on specific Solanaceae lineages. The ages of clades in the Solanaceae phylogeny^42^ were minimum age estimates because Solanaceae fossils known at that time were placed conservatively at the stem ages of lineages with which they shared morphological synapomorphies, and because fossils in general can only provide minimum age estimates for clades. Therefore, using those minimum Solanaceae ages as maximum calibrations for Ithomiini lineages may strongly underestimate the ages of the butterfly lineages, especially when using mean or median age estimates instead of older bounds. Here, we took advantage of a recent calibration of the family-level phylogeny of the angiosperms based on 151 fossils^43^ to recalibrate the Solanaceae phylogeny using the inferred stem age of Solanaceae, i.e., 66.6 mya, as a calibration point in a Bayesian framework (Supplementary Methods S1.2). Due to the recent description of a 52.2 my old Solanaceae fossil placed in the extant genus *Physalis*^55^ that could have dramatic consequences on the age estimates of Solanaceae and Angiosperms as a whole, we also ran an analysis without calibration based on host-plant ages. Indeed, much older ages of Solanaceae lineages as those implied by this discovery would have hardly any impact on the ages of Ithomiini, and removing host-plant derived calibrations is therefore a conservative way of testing the influence of older host-plant ages. Since the two calibration strategies yielded almost the same ages (see results) we performed the biogeographic and diversification analyses on the tree calibrated using the combined calibration strategy detailed below. For further discussion on the recently described Solanaceae fossil and its potential effect on host plant age estimates, see Discussion above.

We extracted the new maximum ages of several Solanaceae lineages from our newly calibrated phylogeny in order to provide maximum calibrations for Ithomiini lineages that feed on them^84^. The choice of the host-plant calibrations (Fig. 1) followed that of the genus-level Ithomiini phylogeny study^37^, with the following exceptions. We excluded the *Solanum* calibration, because the two lineages feeding exclusively on *Solanum* (the Mechanitina and the clade comprising the Oleriina, Dircennina, Godyridina, Ithomiina and Napeogeneina) do not have a sister relationship in our and in other Ithomiini phylogenies^27,34^. We also excluded the *Cestrum* calibration (ithomiine subtribe Godyridina, excluding the genera *Veladyris* and *Velamysta*) because this highly diverse genus had a low sampling fraction of 20% in the Solanaceae phylogeny and was not resolved as monophyletic^42^. Finally, for simplicity we excluded the calibrations based on Solanaceae genera *Brunfelsia* and *Lycianthes*, because they apply to young butterfly lineages (the subtribe Methonina and the clade comprising the genera *Oleria* and *Ollantaya*, respectively^37,38^) and would have no effect on the age estimates. A uniform prior was used for host-plant-derived calibrations. The upper boundary of the prior was set to the upper boundary of the 95% credibility interval of the age of the host-plant lineage and the lower boundary of the prior was set to 0 (present) (Fig. 1).

As a second source of calibration points we used age estimates from Wahlberg *et al*.’s^38^ dated Nymphalidae phylogeny. We defined seven secondary calibrations points which were set to a uniform prior bounded by the upper and lower boundaries of the 95% HPD of the ages inferred in the Nymphalidae phylogeny^38^ (Fig. 1).

Sequences of all *Pteronymia* specimens were combined into a consensus sequence for each species to maximize sequence coverage for each species^34^. We used PartitionFinder 1.1.1^81^ to select the best partition scheme applying to this new dataset, where only the models implemented in BEAST were tested (see Supplementary Table S5 online). A time-tree was generated in BEAST 1.7.5^44^ under an uncorrelated lognormal relaxed clock using the same outgroups as previously, and the dating procedure described above. To select the tree prior (Yule versus Birth-Death), we ran analyses with each type of prior and used Stepping Stone sampling^85^ to estimate marginal likelihood (MLE). This method is not implemented in BEAST 1.7.5, thus we performed this analysis on BEAST 1.8.2 on the molecular dataset only (BEAST 1.8.2 cannot handle morphological characters). The MLE were then used to compute Bayes Factors (BF), which supported the Yule model (BF=4.44). We also ran analyses on the total evidence on BEAST 1.7.5 under the Yule and the Birth-Death model to compare age estimates. Both types of analyses yielded virtually identical ages (regression of ages under Birth-Death (BD) on ages under Yule (Y): BD = 0.9985*Y, r^2^=0.999). Analyses with and without morphological data also produced very similar ages (regression of ages with molecular data only (M) on ages with the combined evidence (C): M = 0.9892*C, r^2^=0.997). Analyses were run for 100 million generations each on the Cipres server^82^ and on a desktop computer depending on the version of BEAST, and trees and parameters were sampled every 100,000 generation. After checking for parameter ESS, a 10% burnin was applied to the posterior distribution and the maximum clade credibility tree with median node ages (hereafter, MCC tree) was computed using TreeAnnotator 1.6.2^44^. Given the results on the tree prior (Yule versus Birth-Death, see above), subsequent analyses were performed on the trees generated under a Yule prior. Because many nodes of the phylogeny had moderate or poor support, we conducted most of the analyses outlined below on the MCC tree and on a random subset of 100 trees extracted from the post burnin posterior distribution of trees obtained from the BEAST run.

### Spatial patterns of diversification

Geographic distribution and elevational range for all extant species were obtained from our own records, museum collections and collaborators (Table 1, Supplementary Fig. S5). The biogeographic history of *Pteronymia* was estimated using the Maximum Likelihood Dispersal-Extinction-Cladogenesis (DEC) model^86^ on the MCC tree, and its extension for multiple trees (Statistical DEC, or S-DEC) on the 100 trees extracted from the posterior distribution, using the software RASP 2.1^45^. We defined nine biogeographical areas (Fig. 2): Central America (A), Western lowlands (B), Slopes of the Western/Central Cordillera of Ecuador and Colombia (C), Central Andes (D), Slopes of the Eastern Cordillera of Colombia and Venezuela (E), Upper Amazon (F), Lower Amazon (G), Atlantic Forest (H), and Guiana Shield (I), and the maximum number of ancestral areas in our analyses was set to four reflecting the maximum number of areas occupied by extant species. We implemented three time slices (11-8, 8-5, 5-0 million years ago, Fig. 1) with different dispersal probabilities, which took into consideration major geological events through time^9^.

### Ancestral state reconstruction and evolution of elevational range

Elevational range (elevation range, i.e., boundaries of the elevational interval containing 95% of the records and mean elevation) for all extant species were extracted from the distributional database computed above (Table 1). Evolution of elevational range was investigated on both the MCC tree and the subset of 100 trees as follows. The phylogenetic signal and the tempo of evolution of the mean elevation and the boundaries of the elevational range were assessed by estimating simultaneously the values of the λ and δ scaling parameters^46
^ that maximized the likelihood of the data using BayesTraits v2^87^. A λ value of one indicates that the phylogeny correctly represents the trait covariance among species, while a value of 0 indicates that the trait evolution is independent of the phylogeny. A value of λ smaller than one indicates that the phylogeny overestimates the trait covariance among species. A δ value of one means that the trait evolves at a constant pace along the branches of the tree; δ < 1 indicates early changes in the character values followed by a slowing down of the evolution rate, such as that entailed by an adaptive radiation; while δ > 1 indicates accelerated evolution rate and species-specific adaptation. The MCC tree was then rescaled with the corresponding δ and λ values inferred for the mean elevation and the upper and lower boundaries of elevational range, when those differed from 1, such that the evolutionary rate of the elevation attributes on the transformed tree was constant. The resulting trees were used to infer ancestral values of the attributes (assuming a constant evolution rate), using the function *contMap* of the R package *phytools*^88^.

### Temporal patterns of diversification

To investigate the pattern of speciation and extinction rate variations through time and across lineages, we chose not to use BAMM 2.5.0^89^ because of recent criticisms on uninformative priors and biased estimates of diversification rates^90^. Instead, we implemented a two-step procedure. We first used MEDUSA^47^ implemented in the R package *geiger*^91^ to detect shifts on the selection of 100 random trees of the posterior distribution, and on the MCC tree. MEDUSA has also been criticised because of its inflated false discovery rate and biased estimates of diversification rates^92^. To overcome these shortcomings and to implement time-dependent models of diversification, we used the method developed by Morlon et al. (2011)^48^, a maximum likelihood approach that accommodates time dependent birth-death processes and enables to test for rate shifts. We used as a rule that if a shift was present in at least 5% of the trees of the posterior distribution in the MEDUSA analysis, this shift was tested with Morlon et al.(2011)’s method^48^. Shifts detected in less than 5% of the posterior distribution were considered non-significant and not tested with Morlon et al. (2011)’s method^48^. We fitted 6 models of diversification^48^: constant speciation without extinction, time-dependent speciation without extinction, constant speciation with constant extinction, time-dependent speciation and constant extinction, constant speciation and time-dependent extinction, time-dependent speciation and time-dependent extinction. Time-dependency was modelled using an exponential function. Models were compared using AIC scores. The root of the tree was always excluded from the analyses. Our phylogeny included all known species so we set the sampling fraction to 1. All models were fitted on the MCC tree and on the 100 trees sampled from the posterior distribution.

**References**

## Acknowledgements

We thank authorities of Peru, Ecuador, Colombia, Panama and Brazil (ICMBio, SISBIO no. 10802-5) for issuing research and collection permits, as well as many assistants for their help in the field and in curating and databasing museum specimens. We are grateful to Haydon Warren-Gash and Niklas Wahlberg for providing specimens. We thank Christine Bacon and Fabien Condamine for providing constructive comments on this manuscript. ME acknowledges financial support from CNRS (France) and the Leverhulme trust (UK). LDS’s postdoc was funded by an ATIP (CNRS, France) grant awarded to ME. NC was funded by a doctoral grant from the Doctoral School 227 (Sciences de la Nature et de l’Homme: Evolution et Ecologie, France). KW acknowledges funding from NSF (DEB-0639861, DEB-0103746), the National Geographic Society, the Darwin Initiative and the Leverhulme Trust. A.V.L.F. thanks CNPq (fellowships 302585/2011-7 and 303834/2015-3), RedeLep-SISBIOTA-Brasil/CNPq (563332/2010-7), BR-BoL (MCT/CNPq/FNDCT 50/2010) and FAPESP (BIOTA-FAPESP Programs 2011/50225-3, 2012/50260-6 and 2013/50297-0). KLSB acknowledges support by FAPESP (2012/16266-6). Support for components of this work was provided through a collaborative grant, Dimensions US-Biota-São Paulo, supported by the US National Science Foundation (NSF DEB 1241056), National Aeronautics and Space Administration (NASA), and the Fundação de Amparo à Pesquisa do Estado de São Paulo (FAPESP Grant 2012/50260-6). Molecular work was performed at the GenePool (University of Edinburgh, UK), UCL (UK) and the Service of Molecular Systematics UMS2700 of the MNHN (France). Work by SK and TS to construct the original Solanaceae phylogeny was funded by the National Science Foundation (DEB-0316614).

### Author Contributions

D.L.D.S., L.L.M., N.C., K.R.W. and M.E. conceived the study, with contributions from A.V.L.F., G.L., T.S., S.K. and C.D.J. D.L.D.S., N.C., R.M., K.L.S-B., L.M.G.P., A.V.L.F., G.L., M.J., J.M., C.E.G., S.U., C.D.J., K.R.W. and M.E. provided *Pteronymia* specimens and sequences. L.L.M., A.V.L.F. and K.R.W. generated the morphological dataset. T.S. and S.K. provided the Solanaceae dataset. D.L.D.S., N.C., K.L.S-B. and M.E. performed the analyses. D.L.D.S., N.C., K.R.W. and M.E. wrote the paper, with contributions from all co-authors.

### Additional Information

#### Supplementary information

Accompanies this paper at http://www.nature.com/srep

### Competing financial interests

The authors declare no competing financial interests.

## References

1. Myers, N., Mittermeier, R.A., Mittermeier, C.G., da Fonseca, G.A.B. & Kent, J. Biodiversity hotspots for conservation priorities. Nature. 403, 853–858 (2000).

2. Beckman, E.J. & Witt, C.C. Phylogeny and biogeography of the New World siskins and goldfinches: Rapid, recent diversification in the Central Andes. Mol. Phylogenet. Evol. 87, 28–45 (2015).

3. Castroviejo-Fisher, S., Guayasamin, J.M., Gonzalez-Voyer, A. & Vila, C. Neotropical diversification seen through glassfrogs. J. Biogeogr. 41, 66–80 (2014).

4. De-Silva, D.L., Elias, M., Willmott, K., Mallet, J. & Day, J.J. Diversification of clearwing butterflies with the rise of the Andes. J. Biogeogr. 43, 44–58 (2016).

5. Ebel, E.R. et al. Rapid diversification associated with ecological specialization in Neotropical Adelpha butterflies. Mol. Ecol. 24, 2392–2405 (2015).

6. Elias, M. et al. Out of the Andes: patterns of diversification in clearwing butterflies. Mol. Ecol. 18, 1716–1729 (2009).

7. Hughes, C. & Eastwood, R. Island radiation on a continental scale: Exceptional rates of plant diversification after uplift of the Andes. Proc. Natl. Acad. Sci. U.S.A. 103, 10334–10339 (2006).

8. Lagomarsino, L.P., Condamine, F.L., Antonelli, A., Mulch, A. & Davis, C.C. The abiotic and biotic drivers of rapid diversification in Andean bellflowers (Campanulaceae). New Phytologist. 210, 1430–1442 (2016).

9. Hoorn, C. et al. Amazonia Through Time: Andean Uplift, Climate Change, Landscape Evolution, and Biodiversity. Science. 330, 927–931 (2010).

10. Gregory-Wodzicki, K.M. Uplift history of the Central and Northern Andes: A review. Geological Society Of America Bulletin. 112, 1091–1105 (2000).

11. Leier, A., McQuarrie, N., Garzione, C. & Eiler, J. Stable isotope evidence for multiple pulses of rapid surface uplift in the Central Andes, Bolivia. Earth and Planetary Science Letters. 371, 49–58 (2013).

12. Bacon, C.D. et al. Biological evidence supports an early and complex emergence of the Isthmus of Panama. Proc. Natl. Acad. Sci. U.S.A. 112, 6110–6115 (2015).

13. Wesselingh, F.P. et al. Lake Pebas: a palaeoecological reconstruction of a Miocene, long-lived lake complex in western Amazonia. Cainozoic Res. 1, 35–81 (2002).

14. Wesselingh, F.P. & Salo, J.A. Miocene perspective on the evolution of the Amazonian biota. Scripta Geol., 133 (2006). Scripta Geologica. 133, 439–458 (2006).

15. Matos-Maraví, P. Investigating the timing of origin and evolutionary processes shaping regional species diversity: Insights from simulated data and neotropical butterfly diversification rates. Evolution. 70, 1638–1650 (2016).

16. Rull, V. Origins of Biodiversity. Science. 331, 398–399 (2011).

17. Rull, V. Pleistocene speciation is not refuge speciation. J. Biogeogr. 42, 602–604 (2015).

18. Smith, B.T. et al. The drivers of tropical speciation. Nature. 515, 406-+ (2014).

19. Gentry, A.H. Neotropical floristic diversity: Phytogeographical connections between Central and South America, Pleistocene climatic fluctuations or an accident of Andean orogeny?. Annals of the Missouri Botanical Garden. 69, 557–593 (1982).

20. Lamas, G. Ithomiinae in J. B. Heppner, ed. Atlas of Neotropical Lepidoptera. Checklist: Part 4A. Hesperioidea - Papilionoidea., (Association for Tropical Lepidoptera/Scientific Publishers, Gainsville, 2004).

21. Antonelli, A. & Sanmartin, I. Why are there so many plant species in the Neotropics? Taxon. 60, 403–414 (2011).

22. Hughes, C.E., Pennington, R.T. & Antonelli, A. Neotropical Plant Evolution: Assembling the Big Picture. Botanical Journal of the Linnean Society. 171, 1–18 (2013).

23. Luebert, F. & Weigend, M. Phylogenetic insights into Andean plant diversification. Frontiers in Ecology and Evolution. 2(2014).

24. McGuire, J.A. et al. Molecular Phylogenetics and the Diversification of Hummingbirds. Curr. Biol. 24, 910–916 (2014).

25. Willmott, K.R., Hall, J.P.W. & Lamas, G. Systematics of *Hypanartia* (Lepidoptera : Nymphalidae: Nymphalinae), with a test for geographical speciation mechanisms in the Andes. Systematic Entomology. 26, 369–399 (2001).

26. Casner, K.L. & Pyrcz, T.W. Patterns and timing of diversification in a tropical montane butterfly genus, Lymanopoda (Nymphalidae, Satyrinae). Ecography. 33, 251–259 (2010).

27. Chazot, N. et al. mutualistic mimicry and filtering by altitude shape the structure of Andean butterfly communities. Am. Nat. 183, 26–39 (2014).

28. Hall, J.P.W. Montane speciation patterns in *Ithomiola* butterflies (Lepidoptera : Riodinidae): are they consistently moving up in the world? Proc. R. Soc. B. 272, 2457–2466 (2005).

29. Matos-Maravi, P.F., Pena, C., Willmott, K.R., Freitas, A.V.L. & Wahlberg, N. Systematics and evolutionary history of butterflies in the “Taygetis clade” (Nymphalidae: Satyrinae: Euptychiina): Towards a better understanding of Neotropical biogeography. Mol. Phylogenet. Evol. 66, 54–68 (2013).

30. Massardo, D., Fornel, R., Kronforst, M., Goncalves, G.L. & Moreira, G.R.P. Diversification of the silverspot butterflies (Nymphalidae) in the Neotropics inferred from multi-locus DNA sequences. Mol. Phylogenet. Evol. 82, 156–165 (2015).

31. Rosser, N., Phillimore, A.B., Huertas, B., Willmott, K.R. & Mallet, J. Testing historical explanations for gradients in species richness in heliconiine butterflies of tropical America. Biol. J. Linnean Soc. 105, 479–497 (2012).

32. Condamine, F.L., Silva-Brandao, K.L., Kergoat, G.J. & Sperling, F.A.H. Biogeographic and diversification patterns of Neotropical Troidini butterflies (Papilionidae) support a museum model of diversity dynamics for Amazonia. BMC Evol. Biol. 12(2012).

33. Beccaloni, G.W. Ecology, behaviour and natural history of ithomiine butterflies (Lepidoptera: Nymphalidae) and their mimics in Ecuador.. Trop. Lep. 8, 103–124 (1997).

34. Chazot, N. et al. Into the Andes: multiple independent colonizations drive montane diversity in the Neotropical clearwing butterflies Godyridina. Mol. Ecol. 25, 5765–5784 (2016).

35. Jiggins, C.D., Mallarino, R., Willmott, K.R. & Bermingham, E. The phylogeneti1c pattern of speciation and wing pattern change in neotropical Ithomia butterflies (Lepidoptera : Nymphalidae). Evolution. 60, 1454–1466 (2006).

36. Chazot, N. et al. Patterns of species, phylogenetic and mimicry diversity of clearwing butterflies in the Neotropics.. in Biodiversity Conservation and Phylogenetic Systematics (eds. Pellens, R. & Grandcolas, P.) 333–354 (Springer, 2016).

37. Garzón-Orduña, I.J., Silva-Brandao, K.L., Willmott, K.R., Freitas, A.V.L. & Brower, A.V.Z. Incompatible Ages for Clearwing Butterflies Based on Alternative Secondary Calibrations. Syst. Biol. 64, 752–67 (2015).

38. Wahlberg, N. et al. Nymphalid butterflies diversify following near demise at the Cretaceous/Tertiary boundary. Proc. R. Soc. B. 276, 4295–4302 (2009).

39. Neild, A. The Butterflies of Venezuela. Part 2: Nymphalidae II (Acraeinae, Libytheinae, Nymphalinae, Ithomiinae, Morphinae). A comprehensive guide to the identification of adult Nymphalidae, Papilionidae, and Pieridae., 276 (Meridian Publications, London, 2008).

40. Bolaños M. I.A., Zambrano G. G. & Willmott, K.R. Descripción de los estados inmaduros de *Pteronymia zerlina zerlina, P. zerlina machay, P. veia florea y P. medellina* de Colombia y del Ecuador (Lepidoptera: Nymphalidae): Ithomiini.. Tropical Lepidoptera Research. 21, 27–33 (2011).

41. Brown, K.S. & Freitas, A.V.L. Juvenile stages of Ithomiinae : overview and systematics (Lepidoptera: Nymphalidae). Trop. Lep. 5, 9–20 (1994).

42. Särkinen, T., Bohs, L., Olmstead, R.G. & Knapp, S. A phylogenetic framework for evolutionary study of the nightshades (Solanaceae): a dated 1000-tip tree. BMC Evol. Biol. 13(2013).

43. Magallón, S., Gomez-Acevedo, S., Sanchez-Reyes, L.L. & Hernandez-Hernandez, T. A metacalibrated time-tree documents the early rise of flowering plant phylogenetic diversity. New Phytologist. 207, 437–453 (2015).

44. Drummond, A.J., Suchard, M.A., Xie, D. & Rambaut, A. Bayesian Phylogenetics with BEAUti and the BEAST 1.7. Molecular Biology and Evolution. 29, 1969–1973 (2012).

45. Yu, Y., Harris, A.J. & He, X.-J. RASP (Reconstruct Ancestral State in Phylogenies), version 2.0. Available: http://mnh.scu.edu.cn/soft/blog/RASP (2013).

46. Pagel, M. Inferring the historical patterns of biological evolution. Nature. 401, 877–884 (1999).

47. Alfaro, M.E. et al. Nine exceptional radiations plus high turnover explain species diversity in jawed vertebrates. Proc. Natl. Acad. Sci. U.S.A. 106, 13410–13414 (2009).

48. Morlon, H., Parsons, T.L. & Plotkin, J.B. Reconciling molecular phylogenies with the fossil record. Proc. Natl. Acad. Sci. U.S.A. 108, 16327–16332 (2011).

49. De-Silva, D.L. et al. Molecular phylogenetics of the neotropical butterfly subtribe Oleriina (Nymphalidae: Danainae: Ithomiini). Mol. Phylogenet. Evol. 55, 1032–1041 (2010).

50. Mallarino, R., Bermingham, E., Willmott, K.R., Whinnett, A. & Jiggins, C.D. Molecular systematics of the butterfly genus *Ithomia* (Lepidoptera : Ithomiinae): a composite phylogenetic hypothesis based on seven genes. Mol. Phylogenet. Evol. 34, 625–644 (2005).

51. Heliconius_Genome_Consortium. Butterfly genome reveals promiscuous exchange of mimicry adaptations among species. Nature. 487, 94–98 (2012).

52. Martin, S.H. et al. Genome-wide evidence for speciation with gene flow in Heliconius butterflies. Genome Research. 23, 1817–1828 (2013).

53. Mavarez, J. et al. Speciation by hybridization in *Heliconius* butterflies. Nature. 441, 868–871 (2006).

54. Foster, C.S.P. et al. Evaluating the Impact of Genomic Data and Priors on Bayesian Estimates of the Angiosperm Evolutionary Timescale. Syst. Biol. syw086(2016).

55. Wilf, P., Carvalho, M.R., Gandolfo, M.A. & Cúneo, N.R. Eocene lantern fruits from Gondwanan Patagonia and the early origins of Solanaceae. Science. 355, 71–75 (2017).

56. Knapp, S. Tobacco to tomatoes: a phylogenetic perspective on fruit diversity in the Solanaceae. Journal of Experimental Botany. 53, 2001–2022 (2002).

57. He, C.Y. & Saedler, H. Heterotopic expression of MPF2 is the key to the evolution of the Chinese lantern of Physalis, a morphological novelty in Solanaceae. Proc. Natl. Acad. Sci. U.S.A. 102, 5779–5784 (2005).

58. Hu, J.Y. & Saedler, H. Evolution of the inflated calyx syndrome in solanaceae. Molecular Biology and Evolution. 24, 2443–2453 (2007).

59. Khan, M.R., Hu, J.Y., Riss, S., He, C.Y. & Saedler, H. MPF2-Like-A MADS-Box Genes Control the Inflated Calyx Syndrome in Withania (Solanaceae): Roles of Darwinian Selection. Molecular Biology and Evolution. 26, 2463–2473 (2009).

60. Sauquet, H. A practical guide to molecular dating. Comptes Rendus Palevol. 12, 355–367 (2013).

61. Blandin, P. & Purser, B. Evolution and diversification of neotropcial butterflies: insights from the biogeography and phylogeny of the genus *Morpho* Fabricius, 1807 (Nymphalidae: Morphinae), with a review of geodynamics of South America. Tropical Lepidoptera Research. 23, 62–85 (2013).

62. Jørgensen, P.M. et al. Regional patterns of vascular plant diversity and endemism. in Climate Change and Biodiversity in the Tropical Andes. Inter-American Institute for Global Change Research (IAI) and Scientific Committee on Problems of the Environment (SCOPE), 192–203. (eds. Herzog, S.K., Martínez, R., Jørgensen, P.M. & Tiessen, H.) 192–203 (2011).

63. Knapp, S. Assessing patterns of plant endemism in neotropical uplands. Bot. Rev. 68, 22–37 (2002).

64. Chamberlain, N.L., Hill, R.I., Kapan, D.D., Gilbert, L.E. & Kronforst, M.R. Polymorphic Butterfly Reveals the Missing Link in Ecological Speciation. Science. 326, 847–850 (2009).

65. Merrill, R.M. et al. Disruptive ecological selection on a mating cue. Proc. R. Soc. B. 279, 4907–4913 (2012).

66. Jiggins, C.D., Naisbit, R.E., Coe, R.L. & Mallet, J. Reproductive isolation caused by colour pattern mimicry. Nature. 411, 302–305 (2001).

67. Merrill, R.M. et al. Mate preference across the speciation continuum in a clade of mimetic butterflies. Evolution. 65, 1489–1500 (2011).

68. Merrill, R.M., van Schooten, B., Scott, J.A. & Jiggins, C.D. Pervasive genetic associations between traits causing reproductive isolation in Heliconius butterflies. Proc. R. Soc. B. 278, 511–518 (2011).

69. McClure, M. & Elias, M. Ecology, life history, and genetic differentiation in the Neotropical Melinaea (Nymphalidae: ithomiini) butterflies from north-eastern Peru.. Zoological Journal of the Linnean Society. Early view. (2016).

70. Elias, M. et al. Phylogenetic hypothesis, pattern of speciation and evolution of wing pattern in neotropical Napeogenes butterflies (Lepidoptera: Nymphalidae). in 7th International Workshop on the Molecular Biology and Genetics of the Lepidoptera August 20–26, 2006, Orthodox Academy of Crete, Kolympari, Crete, Greece. 52pp., Vol. 7:29 (eds. Iatrou, K. & Couble, P.) 13–14 (Journal of Insect Science, 2007).

71. Antonelli, A., Nylander, J.A.A., Persson, C. & Sanmartin, I. Tracing the impact of the Andean uplift on Neotropical plant evolution. Proc. Natl. Acad. Sci. U.S.A. 106, 9749–9754 (2009).

72. Bloch, J.I. et al. First North American fossil monkey and early Miocene tropical biotic interchange. Nature. 533, 243-+ (2016).

73. Farris, D.W. et al. Fracturing of the Panamanian Isthmus during initial collision with South America. Geology. 39, 1007–1010 (2011).

74. Montes, C. et al. Middle Miocene closure of the Central American Seaway. Science. 348, 226–229 (2015).

75. Sedano, R.E. & Burns, K.J. Are the Northern Andes a species pump for Neotropical birds? Phylogenetics and biogeography of a clade of Neotropical tanagers (Aves: Thraupini). J. Biogeogr. 37, 325–343 (2010).

76. Velazco, P.M. & Patterson, B.D. Diversification of the Yellow-shouldered bats, Genus Sturnira (Chiroptera, Phyllostomidae), in the New World tropics. Mol. Phylogenet. Evol. 68, 683–698 (2013).

77. Goloboff, F., Farris, J.S. & Nixon, K.C. TNT: Tree Analysis using New Technology. Program and documentation, available from the authors, and at http://www.zmuc.dk/public/phylogeny. (2003).

78. Nixon, K.C. Winclada (Beta). Published by the author, Ithaca, NY. (1999).

79. Stamatakis, A., Hoover, P. & Rougemont, J. A Rapid Bootstrap Algorithm for the RAxML Web Servers. Syst. Biol. 57, 758–771 (2008).

80. Ronquist, F. et al. MrBayes 3.2: Efficient Bayesian Phylogenetic Inference and Model Choice Across a Large Model Space. Syst. Biol. 61, 539–542 (2012).

81. Lanfear, R., Calcott, B., Ho, S.Y.W. & Guindon, S. PartitionFinder: Combined Selection of Partitioning Schemes and Substitution Models for Phylogenetic Analyses. Molecular Biology and Evolution. 29, 1695–1701 (2012).

82. Miller, M.A., Pfeiffer, W. & Schwartz, T. Creating the CIPRES Science Gateway for Inference of Large Phylogenetic Trees. . SC10 Workshop on Gateway Computing Environments (GCE10). (2010).

83. Huelsenbeck, J.P., Larget, B. & Alfaro, M.E. Bayesian phylogenetic model selection using reversible jump Markov chain Monte Carlo. Molecular Biology and Evolution. 21, 1123–1133 (2004).

84. Willmott, K.R. & Freitas, A.V.L. Higher-level phylogeny of the Ithomiinae (Lepidoptera : Nymphalidae): classification, patterns of larval hostplant colonization and diversification. Cladistics. 22, 297–368 (2006).

85. Xie, W.G., Lewis, P.O., Fan, Y., Kuo, L. & Chen, M.H. Improving Marginal Likelihood Estimation for Bayesian Phylogenetic Model Selection. Syst. Biol. 60, 150–160 (2011).

86. Ree, R.H. & Smith, S.A. Maximum likelihood inference of geographic range evolution by dispersal, local extinction, and cladogenesis. Syst. Biol. 57, 4–14 (2008).

87. Pagel, M., Meade, A. & Barker, D. Bayesian estimation of ancestral character states on phylogenies. Syst. Biol. 53, 673–684 (2004).

88. Revell, L.J. phytools: an R package for phylogenetic comparative biology (and other things). Methods in Ecology and Evolution. 3, 217–223 (2012).

89. Rabosky, D.L. Automatic Detection of Key Innovations, Rate Shifts, and Diversity-Dependence on Phylogenetic Trees. PLoS ONE. 9(2014).

90. Moore, B.R., Hohna, S., May, M.R., Rannala, B. & Huelsenbeck, J.P. Critically evaluating the theory and performance of Bayesian analysis of macroevolutionary mixtures. Proc. Natl. wAcad. Sci. U.S.A. 113, 9569–9574 (2016).

91. Pennell, M.W. et al. geiger v2.0: an expanded suite of methods for fitting macroevolutionary models to phylogenetic trees. Bioinformatics. 30, 2216–2218 (2014).

92. May, M.R. & Moore, B.R. How Well Can We Detect Lineage-Specific Diversification-Rate Shifts? A Simulation Study of Sequential AIC Methods. Syst. Biol. 65, 1076–1084 (2016).

